# Isoform-Resolved Genetic Architecture of Epilepsy and SUDEP Reveals Divergent Brain and Heart Channelopathy Signatures

**DOI:** 10.64898/2026.03.31.715736

**Authors:** Binte Zehra, Shuhd BinEshaq, Muhammad Faizan, Mariam Eldesouky, Nidhina Vinod, Nesrin Mohamed, Aswathy Vijayakumar, Inna Aleksandrova, Richa Tambi, Solima Sabeel, Dia Advani, Amiruddin Hashmi, Shaiban Al-Shaibani, Mohamed Almarri, Nasna Nassir, Sumaya Almansoori, Stefan S. Du Plessis, Mohammed Uddin, Bakhrom K. Berdiev

**Author notes:** Corresponding Authors: Bakhrom K. Berdiev, Mohammed Uddin. These authors have contributed equally to this work (Binte Zehra, Shuhd BinEshaq).

## Abstract

Sudden unexpected death in epilepsy (SUDEP) is the most devastating complication of epilepsy, yet the molecular features distinguishing individuals at risk remain poorly defined. Although epilepsy and SUDEP share substantial genetic overlap, fatal outcomes may arise when shared risk genes are differentially deployed across neuronal and cardiac systems. Here, we identify tissue- and isoform-level regulation as a key determinant of divergence between epilepsy and SUDEP risk. We performed a large-scale integrated analysis of genetic variants reported in epilepsy and SUDEP across 419 sequencing-based studies encompassing 35,659 individuals, and quantified gene-level burden using a Bayesian Poisson–Gamma rate ratio framework. This analysis revealed preferential enrichment of genes related to cardiac electrophysiology and contractile function in SUDEP, whereas epilepsy was dominated by genes involved in neuronal excitability and synaptic signaling. To determine how shared genetic loci are deployed across tissues, we integrated GTEx-based tissue expression profiles with long-read single-cell transcriptomic datasets from human heart and brain to resolve isoform-level expression patterns. These analyses revealed pronounced tissue-specific transcript architectures. Cardiac-associated genes, including *HCN4, KCNH2, KCNE1, MYH6, MYO18B*, and *ATP1A2,* showed heart-restricted isoform expression, whereas neuronal genes such as *ADGRV1, CACNA1A, GRIN2B, HCN1, HCN2, KCNA1, SCN1A, SCN2A,* and *SCN8A*. Importantly, several shared genes exhibited tissue-partitioned isoform expression, with distinct transcript repertoires in heart and brain, particularly across pathways related to ion transport, signaling, metabolism, and structural organization. Consistent patterns were observed in iPSC-derived cardiomyocytes and neurons, indicating that lineage-dependent deployment of shared genes is preserved in controlled systems. Together, these findings suggest that tissue-specific isoform regulation provides a mechanistic basis linking shared epilepsy genetics to SUDEP susceptibility, whereby the same genetic loci contribute to neuronal dysfunction in epilepsy and to cardiac vulnerability in SUDEP. This positions SUDEP as a neuro–cardiac interface disorder shaped by isoform-level regulatory divergence.

## Introduction

Epilepsy is a neurological disorder characterized by recurrent seizures resulting from abnormal and excessive electrical activity in the brain (Thijs et al., 2019). With a global prevalence of approximately 7.6 per 1,000 individuals, the condition affects millions of people worldwide and is particularly common among children and adolescents (Tong et al., 2023). Beyond the burden of recurrent seizures, epilepsy is also associated with a substantial risk of premature mortality. Epidemiological studies report that the standardized mortality ratio (SMR) in individuals with epilepsy can be up to 24 times higher than in the general population (Mbizvo et al., 2019; Trinka et al., 2023). Among the most serious contributors to this excess mortality is Sudden Unexpected Death in Epilepsy (SUDEP), which represents one of the leading and most devastating causes of death among individuals with epilepsy (Stewart, 2018).

SUDEP is a multifactorial condition involving dysfunction across neurological, respiratory, and cardiac systems and occurs at an estimated rate of approximately 1 per 1,000 individuals with epilepsy each year (Cihan et al., 2020). In individuals with severe, refractory epilepsy, however, the risk is markedly higher, with reported incidences ranging from 1 in 150 to 1 in 300 individuals per year (O’Neal et al., 2022). Clinical observations suggest that SUDEP frequently follows a generalized tonic–clonic seizure, during which postictal arousal dysfunction can lead to impaired breathing and severe cardiac arrhythmias, including bradycardia or asystole (Muccioli et al., 2024; Ryvlin et al., 2013; Van der Lende et al., 2016). Established risk factors include uncontrolled tonic–clonic seizures, nocturnal seizures, poor adherence to anti-epileptic medications, and genetic susceptibility involving variants in genes associated with neuronal excitability or cardiac electrophysiology (Sveinsson et al., 2020; Uddin et al., 2017).

Genetic predisposition is increasingly recognized as an important contributor to SUDEP risk. Post-mortem and sequencing-based studies have identified pathogenic variants in several epilepsy- and cardiac-related genes in SUDEP cases (Sahly et al., 2022). For example, variants in sodium channel–encoding genes expressed in both brain and heart, including *SCN1A, SCN1B, SCN2A,* and *SCN5A*, have been associated with increased susceptibility to SUDEP (Sahly et al., 2022). Additional genes involved in cardiac excitability and repolarization, such as *HCN4* and *SCN5A,* which are also implicated in cardiac arrhythmias and Long QT syndrome, have likewise been screened for pathogenic variants in SUDEP cohorts (Bagnall et al., 2016). Importantly, emerging evidence suggests that SUDEP risk may not arise from a single causal mutation but rather from additive effects of multiple variants across genes expressed in neuronal and cardiac tissues, potentially creating combined vulnerabilities across organ systems (Whitney et al., 2023). Despite these advances, the genetic architecture of SUDEP remains incompletely understood, and the mechanisms through which genetic variants influence the transition from epilepsy to sudden death remain largely unresolved.

In this study, we hypothesized that although epilepsy and SUDEP share a common pool of genetic variants, these variants are differentially deployed across tissues, leading to distinct molecular mechanisms underlying each phenotype. We further proposed that genotype–phenotype relationships diverge due to tissue-specific gene regulation and isoform-level expression differences between neuronal and cardiac systems. In this framework, mutations within the same gene may produce distinct functional outcomes depending on the cellular context in which the gene is expressed, effectively partitioning neuronal excitability from cardiac conduction and contractile programs. To test this hypothesis, we performed a comprehensive comparative analysis of genetic variants reported in SUDEP and epilepsy across publicly available sequencing studies. Variants were systematically classified by type, origin, predicted functional effect, and pathogenicity to identify recurrent SUDEP-associated variants. Gene- and domain-level mapping was used to localize variants within protein structures. In parallel, integration of publicly available long-read single-cell isoform datasets from adult human left ventricle and prefrontal cortex (PFC), together with iPSC-derived neurons and cardiomyocytes, enabled characterization of cell-type–specific and isoform-resolved expression patterns of genes shared between epilepsy and SUDEP. Collectively, this integrative framework allowed us to dissect the genetic and regulatory architecture underlying both phenotypes and to provide mechanistic insight into how shared genetic risk can drive epilepsy through neuronal dysfunction while increasing susceptibility to sudden death through cardiac-specific transcript dysregulation.

## Methods

### Data Collection

Genomic variant data were compiled from previously published studies of epilepsy and SUDEP identified through structured searches of PubMed and Google Scholar. The final search was conducted in July 2025 and included all peer-reviewed articles published up to that date in English. Search queries incorporated combinations of Medical Subject Headings (MeSH) terms and keywords, including “whole genome sequencing,” “whole exome sequencing,” “epilepsy cohort variants,” “SUDEP variants,” and “SUDEP sequencing,” using Boolean operators (AND/OR) to optimize retrieval. Search strategies were adapted to the syntax and functionality of each database. A formal PRISMA framework was not applied, as this study represents an integrative analysis rather than a systematic review.

The literature search and study selection were performed by S.B., M.E., and N.V. Studies were included if they reported genomic or exonic sequencing data with accessible variant-level information and provided sufficient genomic annotation, including chromosomal coordinates, gene symbols, or transcript identifiers. Studies describing non-coding variants outside exonic or canonical splice regions, duplicate reports, or publications lacking sufficient genomic detail were excluded (Supplementary Files). The included datasets encompassed epilepsy-related phenotypes such as developmental and epileptic encephalopathies, focal and generalized epilepsies, and febrile or infantile seizures, as well as SUDEP cases with available exome or genome variant data.

Variant-level information was systematically extracted from the selected publications for downstream analysis (Supplementary Files). Extracted fields included genomic coordinates (hg38 reference genome), gene symbol, reference and alternative alleles, codon and amino acid changes, reference SNP identifier (rsID), variant type, variant origin (germline or *de novo*), and predicted functional consequence. Additional metadata such as PubMed identifiers (PMIDs) and sequencing platform information were also recorded.

Variants originally reported in the hg19 genome build were converted to hg38 using the UCSC LiftOver tool (https://genome-euro.ucsc.edu/cgi-bin/hgLiftOver) to standardize genomic coordinates across datasets. Following harmonization, descriptive analyses were performed to examine gene frequency distributions and phenotype associations.

### Variant Annotation

Variants were annotated using ANNOVAR (ANNOtate VARiation) and further evaluated using the GenomeArc Horizon™ platform (Eldesouky et al., 2026) (v2.0). Variants were first filtered against population allele frequencies in gnomAD (Genome Aggregation Database, v4.1), and only rare variants (gnomAD AF < 0.001) were retained for analysis. Variants located outside exonic or canonical splice regions were excluded. To prioritize potentially deleterious variants for downstream interpretation, variants with a Combined Annotation Dependent Depletion (CADD) score ≥ 20 were highlighted for identifying genes shared between epilepsy and SUDEP cohorts (Supplementary Files). Variants classified as benign or likely benign in ClinVar were excluded from downstream analyses. Variant pathogenicity was evaluated using the American College of Medical Genetics and Genomics (ACMG) guidelines, which classify variants into five categories: benign, likely benign, variant of uncertain significance (VUS), likely pathogenic, and pathogenic. Additional *in silico* prediction scores were incorporated to support pathogenicity assessment, including SIFT (Damaging), PolyPhen-2 (Probably Damaging), and REVEL (>0.5 for missense variants). Genotype–phenotype annotations available in ClinVar were also integrated into the evaluation process. Finally, filtered variants were mapped to their corresponding genes and protein domains using the ProteinPaint visualization platform (https://pecan.stjude.org/), enabling domain-level interpretation of variant distribution across proteins.

### Mutation Rate Estimation

To delineate the shared and distinct genetic architectures underlying SUDEP and epilepsy, we quantified gene-specific differences in mutational burden between the two cohorts. A Bayesian Poisson–Gamma rate ratio framework was applied to estimate posterior rate ratios (RRs) while accounting for disparities in cohort size and the sparse nature of rare variant counts. For each gene, the aggregated variant burden was calculated separately for the SUDEP and epilepsy cohorts after applying the initial filtering criteria (*see Variant Annotation*).

Variants were assigned functional weights proportional to their predicted deleteriousness, and the weighted counts were summed for each gene to obtain expected mutational burdens *K_g,s_* and *K_g,e_* for the SUDEP and epilepsy cohorts, respectively. These gene-level counts were modeled under a Poisson likelihood with Gamma priors on the underlying mutation rates, enabling estimation of posterior rate ratios that reflect the relative enrichment of variants in SUDEP compared with epilepsy.

Bayesian Poisson–Gamma Model:

*Gene-level counts were modelled using Poisson likelihoods with conjugate Gamma priors:*

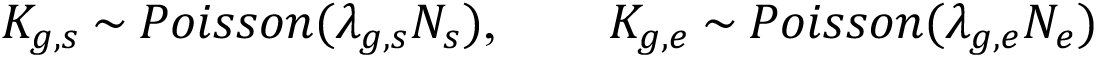

where *N_s_* = 698 and *N_e_* = 21,943 represent the number of SUDEP and epilepsy cases, respectively. To regularize sparse or zero-inflated data, we used weakly informative proper priors *λ*_*g,s*_(*SUDEP*) & *λ_g,e_* (*epilepsy*) ∼ Gamma (1, 1) yielding posteriors:

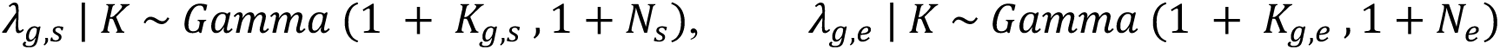

from which we estimated the posterior rate ratio using 200,000 Monte Carlo samples per gene, as:

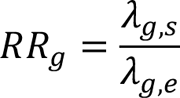

The posterior median and 95% credible interval (Crl) were reported, together with the posterior probability *P*(*RR_g_* > 1). False discovery rates (FDR) were estimated by applying the Benjamini–Hochberg procedure to posterior tail probabilities (*p* = 1 − *P*(*RR_g_* > 1)). Genes with RR > 1 and FDR < 0.05 were classified as *SUDEP-enriched*, whereas those with RR < 1 and FDR < 0.05 were considered *epilepsy-enriched*.

### Pathway Enrichment Analysis and Network Visualization

Pathway enrichment analysis was conducted to explore the molecular pathways associated with the identified epilepsy and SUDEP genes. This analysis was performed using the GeneOverlap package in R in combination with Cytoscape. The parameters for enrichment included a minimum gene overlap of five, a false discovery rate (FDR) threshold of 0.01, and a cutoff p-value of 0.001. Both the Kyoto Encyclopedia of Genes and Genomes (KEGG) and Gene Ontology (GO) databases were used as references for pathway enrichment. The resulting networks were visualized in Cytoscape using the AutoAnnotate and EnrichmentMap applications. In the final pathway maps, node color represents the level of statistical significance (*p*-value), while node size corresponds to the odds ratio.

### iPSC Culture and Lineage Differentiation

Human induced pluripotent stem cells (iPSCs) purchased from Gibco, Life Technologies, were used for both neuronal and cardiomyocyte differentiation. iPSCs were cultured on precoated plates with 10 μg/ml Rh-laminin-521 (Life Technologies) and maintained in Essential 8™ (E8) medium supplemented with 10 μM Revita™ at 37 °C in a 5% CO₂ incubator. On the following day, the medium was replaced with fresh E8, and cells were maintained until they reached 80–90% confluency.

#### Neuronal Differentiation

For neuronal generation, confluent iPSCs were reprogrammed into neural progenitor cells (NPCs) and subsequently differentiated into neurons following our previously established protocol (Zehra et al., 2024). Briefly, 5 × 10⁵ cells were seeded in E8 medium containing 10 μM Revita on 6-well plates coated with laminin and incubated overnight. The next day, the medium was replaced with neural induction medium (Neurobasal medium + 1% Neural Induction Supplement, Life Technologies). To generate NPCs, the medium was refreshed every other day until day 7. On day 8, Neural Expansion Medium (0.5× Neurobasal medium, 0.5× KnockOut™ DMEM/F12, 1× Neural Induction Supplement) was added, and cells were maintained until ∼90% confluency. On day 10, NPCs were harvested using TrypLE Express (Life Technologies), cryopreserved, and later used for neuronal differentiation.

For neuronal differentiation, 1 × 10⁶ NPCs were seeded on poly-D-lysine/Rh-laminin–coated plates in Neural Expansion Medium supplemented with 10 μM ROCK inhibitor. After overnight incubation at 37 °C, the medium was replaced with neuronal differentiation medium (0.5× Neurobasal medium, 0.5× DMEM/F-12, 1× B27 supplement, 0.5× N2 supplement, 125 mM GlutaMAX, and 10 ng/ml bFGF). Cells were maintained in neuronal differentiation medium with media changes every other day. Further experiments were performed using post-D30 iPSC-derived neurons.

#### Cardiomyocyte Differentiation

For cardiomyocyte differentiation, 5 × 10⁵ iPSCs were seeded in E8 medium supplemented with 10 μM Revita on Geltrex™-coated 6-well plates (Life Technologies) and incubated overnight. The next day, the medium was replaced with Cardiomyocyte Induction Medium (RPMI 1640 containing 1× B27 supplement without insulin) and incubated for 24 hours. After mesoderm induction, Cardiomyocyte A Medium (Cardiomyocyte Induction Medium supplemented with 6 µM CHIR99021) was added, and the cells were incubated for 48 hours. The medium was then changed to Cardiomyocyte B Medium (Cardiomyocyte Induction Medium containing 2.5 µM IWP-2) to generate cardiac progenitor cells, followed by incubation for another 48 hours. Subsequently, the medium was replaced with Cardiomyocyte Maintenance Medium (RPMI 1640 supplemented with 1× B27) to promote maturation. Cells were maintained in this medium until day 15, with medium changes performed every other day. The beating iPSC-derived cardiomyocytes were then harvested and used for downstream experiments.

### Tissue-Specific Gene Expression Analysis Using GTEx

To evaluate the tissue-specific expression of genes implicated in epilepsy and SUDEP, we utilized bulk RNA-seq data from the Genotype-Tissue Expression (GTEx) Project, Analysis Release V10. The dataset was downloaded from the GTEx portal (https://gtexportal.org/), which provides transcript-per-million (TPM) normalized expression values generated using RNA-SeQC v2.4.2 across multiple human tissues.

From this dataset, expression profiles corresponding to brain and cardiac tissues were extracted. Brain samples included thirteen regions: frontal cortex (BA9), cerebellar hemisphere, substantia nigra, anterior cingulate cortex (BA24), amygdala, caudate (basal ganglia), nucleus accumbens (basal ganglia), putamen (basal ganglia), cortex, hypothalamus, cerebellum, spinal cord (cervical c-1), and hippocampus. Cardiac samples included the left ventricle and atrial appendage. Only protein-coding genes were retained, and TPM values were log₂-transformed after adding a pseudo-count of 0.1, that is (log₂ [TPM + 0.1]) to stabilize variance. For each gene, mean expression across all brain samples and across all heart samples was computed to obtain representative values per tissue group. The log₂ fold-change (Heart/Brain) was calculated for each gene to determine tissue bias in expression:

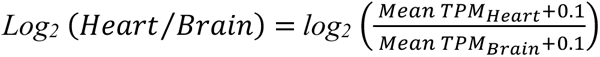

Visualization and statistical processing were performed in Python (v3.10.3) using Pandas, NumPy, and Matplotlib/Seaborn libraries. Bar-coded log₂ fold-change heatmap was used to summarize tissue bias for all genes of interest.

### Long Read Single-Cell mRNA Sequencing

#### In-House Long-Read scRNA-seq Data

Differentiated iPSCs-neurons and iPSCs-cardiomyocytes were collected into single-cell suspension using TrypLE Express and washed with 0.4% BSA prepared in PBS (pH 7.0), followed by filtration through a 70 µm cell strainer to ensure single-cell purity. Cell viability and concentration were assessed using AO/PI fluorescent dye, and only suspensions with 85–90% viable cells and an adjusted density of 1,200 cells/µL were used for Gel Bead-in-Emulsion (GEM) generation on the 10x Chromium platform, targeting a recovery of approximately 10,000 cells per sample. Full-length barcoded cDNA was captured using Dynabeads MyOne SILANE (10x Genomics) and amplified through 13 PCR cycles. cDNA fragment size and concentration were assessed using Agilent TapeStation and Qubit Fluorometric Assay, respectively. A total of 10 ng of amplified cDNA was used to prepare long-read sequencing libraries following the Oxford Nanopore Technologies (ONT) single-cell cDNA sequencing protocol.

For pre-pull-down PCR, 10 µM biotinylated forward primer (5’- /5Biosg/CAGCACTTGCCTGTCGCTCTATCTTCCTACACGACGCTCTTCCGATCT-3’) and partial_TSO_defined reverse primer (5’-CAGCTTTCTGTTGGTGCTGATATTGCAAGCAGTGGTATCAACGCAGAG-3’) were used with LongAmp Hot Start Taq polymerase. Thermal cycling conditions were: initial denaturation at 94°C for 3 min, followed by 30 cycles of denaturation (94°C, 30 s), annealing (66°C → 58°C, ramp down 0.2°C/s for 90 s), and extension (65°C, 6 min), ending with a final extension at 65°C for 10 min. Biotinylated cDNA was purified using AMPure XP beads and captured on M280 streptavidin beads. After two washes with binding buffer (10 mM Tris-HCl pH 7.5, 2 M NaCl, 1 mM EDTA), bead–cDNA conjugates were resuspended in 10 mM Tris-HCl and eluted in nuclease-free water. The purified cDNA was further amplified using the cPRM primer and LongAmp Hot Start Taq polymerase under the following conditions: initial denaturation at 94°C for 3 min, followed by 15 cycles of denaturation (94°C, 15 s), annealing (62°C, 15 s), and extension (65°C, 6 min), with a final extension at 65°C for 10 min. The final full-length barcoded cDNA library was purified with AMPure XP beads, and Rapid Adapter T-ligation (PCS-111) was performed. Approximately 50 fmol of library was loaded onto a PromethION R9.4.1 (FLO-PRO002) flow cell with the corresponding sequencing buffer and Loading Beads II (LBII) to acquire the sequencing data.

Raw FASTQ files were processed using the epi2me-labs/wf-single-cell pipeline (Oxford Nanopore Technologies, v3.3.1). Per-read quality and length statistics were generated using Fastcat v0.3.3 and Bamstat v2.0.1. Adapter trimming and read quality filtering were performed to ensure the inclusion of high-quality, full-length reads. Cell barcodes and unique molecular identifiers (UMIs) were extracted and error-corrected following the ONT single-cell specification. Reads were aligned to the GRCh38 human reference genome (Ensembl v110) using Minimap2 v2.28 with strand-specific alignment enabled. Gene- and transcript-level expression matrices were subsequently generated and merged for downstream analysis. For quality control, the Median Absolute Deviation (MAD) method was applied to remove low-quality or damaged cells, using a threshold defined as the median value plus three times the MAD for nFeature_RNA, nCount_RNA, and mitochondrial transcript fraction. Cells with fewer than three detected genes or high mitochondrial content were excluded. Downstream normalization, scaling, and dimensionality reduction were performed using Scanpy v1.10.2 in Python v3.10.3. Principal Component Analysis (PCA) was conducted on the top 2,000 highly variable genes, followed by Uniform Manifold Approximation and Projection (UMAP) for low-dimensional visualization of transcriptionally distinct clusters. Clustering was performed using the Leiden algorithm based on the top 20 principal components, with the resolution parameter empirically optimized to achieve stable and biologically distinct cluster separations.

Differentially expressed genes (DEGs) were identified using the Wilcoxon rank-sum test with an adjusted p-value threshold of ≤ 0.05. A gene was defined as a cluster marker if expressed in more than 20% of cells within a cluster and showed significantly higher expression compared to all other clusters. For cell-type annotation, canonical marker genes were curated from recent literature on neuronal and cardiomyocyte differentiation, and expression patterns were visualized using dot plots, violin plots, and feature maps. All visualizations were generated in Scanpy using default settings, and final figures were refined in Seaborn v0.13.2 and Matplotlib v3.8.4 for visual clarity and consistency.

#### Publicly Available Long-Read scRNA-seq Data

Publicly available long-read single-cell RNA sequencing datasets were obtained from the Gene Expression Omnibus (GEO) and accessed through the NCBI Sequence Read Archive (SRA). Cardiac long-read datasets were derived from accession GSE288222 (Pan et al., 2025), comprising non-failing donor human heart samples, whereas brain long-read datasets were obtained from accession GSE178175 (Hardwick et al., 2022), consisting of human cortical tissue samples.

For the cardiac dataset (GSE288222), six non-diseased donor heart samples were analyzed: GSM8761493 (ND178), GSM8761494 (ND182), GSM8761495 (ND202), GSM8761496 (ND236), GSM8761497 (ND245), and GSM8761498 (ND248). For the brain dataset (GSE178175), two Oxford Nanopore Technologies (ONT) long-read single-cell RNA sequencing samples from the human frontal cortex were analyzed: GSM5381317 (FCtx1_ONT) and GSM5381318 (FCtx2_ONT).

Raw sequencing data were downloaded using the SRA Toolkit. Archived files were retrieved with prefetch, converted to FASTQ format using fasterq-dump, and compressed using parallel gzip compression (pigz). Raw FASTQ files were processed using the Oxford Nanopore Technologies epi2me-labs/wf-single-cell Nextflow pipeline (v3.3.1). Samples were analyzed using the 3′ single-cell kit configuration (–kit 3prime:v3) with an expected cell number of 10,000 (–expected_cells 10000) and the prebuilt GRCh38 resource bundle (gex-GRCh38-2024-A) corresponding to Ensembl release v110. Within the workflow, input reads were concatenated and summarized using fastcat/bamstats (v0.22.0). Adapter sequences were identified and trimmed, and cell barcodes and unique molecular identifiers (UMIs) were extracted and corrected. Reads were aligned to the GRCh38 reference genome using minimap2 (v2.24) within the workflow. Gene- and transcript-level count matrices were generated, including transcriptome assembly with StringTie (v2.2.3) and annotation using gffcompare.

Gene-level and transcript-level count matrices were produced through the ONT workflow, preserving isoform-resolved transcript assignments. Expression matrices from individual samples were merged separately for the heart and brain datasets. Quality control and downstream analyses were performed using Scanpy (v1.11.5) in Python. Cells with fewer than 200 detected genes were removed, and genes expressed in fewer than three cells were excluded. Additional filtering removed outlier cells based on median absolute deviation (MAD) thresholds applied to log-transformed total UMI counts and the number of detected genes. Cells with high mitochondrial content were filtered using a combined criterion of 3 MADs above the median mitochondrial fraction or an absolute mitochondrial read fraction cutoff (≤25% for cardiac samples and ≤2% for brain samples), reflecting tissue- and dataset-specific mitochondrial content distributions. Library-size normalization was performed independently for each dataset by scaling each cell to 10,000 counts, followed by log transformation.

To correct for donor- and batch-related variability within each tissue, dataset integration was performed using Harmony, with sample identity specified as the batch variable. Harmony was applied to the PCA embeddings to generate batch-corrected low-dimensional representations. For downstream analyses, tissue-specific numbers of Harmony-corrected principal components were used to construct k-nearest neighbor graphs. 40 Harmony-corrected PCs were used for the cardiac dataset, whereas 25 PCs were used for the brain dataset. Uniform Manifold Approximation and Projection (UMAP) was computed from the resulting neighbor graphs, and clustering was performed using the Leiden algorithm. For the cardiac dataset, Leiden clustering was initially performed at resolution 0.1 and subsequently refined at resolution 0.3 to resolve specific cell populations prior to downstream analyses. For the brain dataset, Leiden clustering was performed at resolution 0.1 for cell-type annotation.

Cell-type annotation was conducted using curated lineage-specific marker gene sets. In cardiac tissue, cell populations were defined using canonical markers for cardiomyocytes (*TTN, TECRL, MYH6)*, endothelial cells (*VWF, BTNL9, NOSTRIN, CD34, ADGRL4*), cardiac fibroblasts (*COL4A4, ABCA6, FBLN1, FBN1*), pericytes (*EGFLAM, RGS5, GUCY1A2, PDGFRB*), smooth muscle cells (*MYH11, LGR6*), macrophages/myeloid cells (*CD163, STAB1, RBM47, LYZ, CD86, CD14*), endocardial cells (*SMOC1, EMCN, NPR3*), neural-associated cells (XKR4, PTPRZ1, KCNH8), T cells (*CD247, THEMIS, RUNX3*), and lymphatic endothelial cells (*TBX1, PIEZO2*).

For cortical brain tissue, cell types were annotated using established neural lineage markers, including excitatory neurons (*SLC17A7, SLC17A6, CAMK2A, GRIN1, SATB2*), inhibitory neurons (*GAD1, GAD2, SLC6A1, SLC32A1, DLX1, DLX2*), astrocytes (*ALDH1L1, GFAP, SLC1A3, AQP4, S100B*), oligodendrocytes (*MBP, PLP1, MOG, MAG, MOBP*), microglia (*P2RY12, CX3CR1, TMEM119, C1QA, TYROBP*), oligodendrocyte precursor cells (*PDGFRA, CSPG4*), and endothelial cells (*PECAM1, KDR, VWF, CLDN5, FLT1*).

#### Comparative Analysis of Tissue-Specific Isoform Expression

Transcript-level expression matrices, which provide full-length read–based counts corresponding to reconstructed transcript isoforms, were used directly for downstream analyses to evaluate relative isoform expression patterns across tissues. To ensure robustness and minimize low-frequency transcriptional noise, transcripts were considered expressed only if detected in at least 100 cells. In addition, only protein-coding transcripts were included based on Ensembl biotype annotation, and transcripts supported by fewer than 10 UMI counts were excluded to reduce spurious signal. Filtered isoforms were then compared between cardiac (left ventricle) and brain (prefrontal cortex) datasets to evaluate tissue-specific isoform expression. Differences in isoform deployment were assessed based on consistent tissue enrichment and relative abundance differences across datasets, and isoforms were considered differentially used when they demonstrated reproducible enrichment in one tissue compared to the other. Analyses were restricted to robust, high-confidence isoforms exhibiting consistent and biologically meaningful tissue-specific patterns.

## Results

We performed a comprehensive literature search in PubMed and Google Scholar to identify original research studies reporting whole-genome sequencing (WGS) and/or whole-exome sequencing (WES) in epilepsy and SUDEP. This search yielded 510 primary studies. After removing duplicates and excluding articles without sequencing data or with insufficient variant annotation, we curated a final set of 419 studies (Fig. 1; Supplementary Tables 1–2). Collectively, these studies contributed genomic data from 34,421 epilepsy cases and 1,238 SUDEP cases, implicating 2,753 genes in epilepsy and 896 genes in SUDEP (Figure 1).

**Figure 1.**
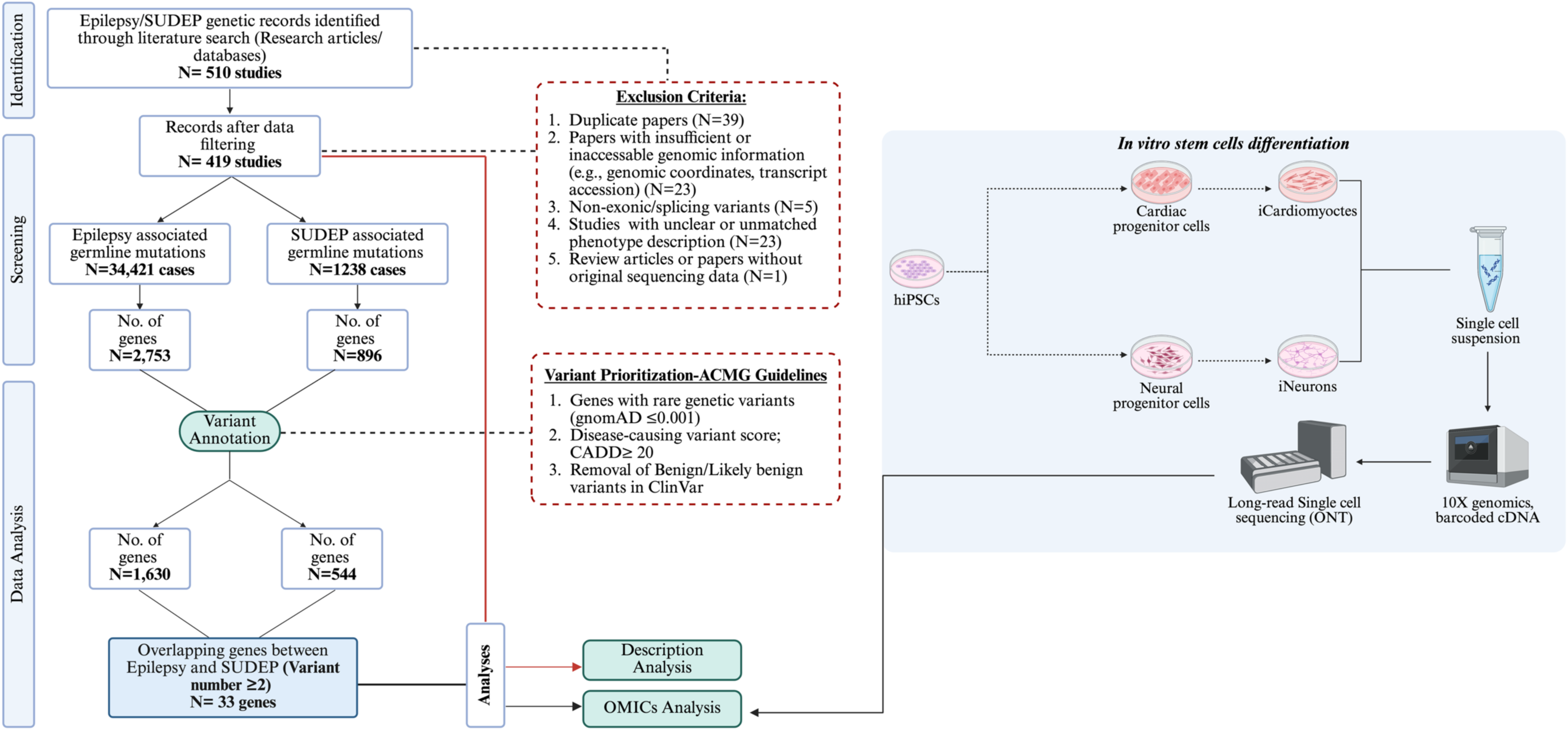
Study workflow and variant prioritization. Schematic overview of study design, literature-based data collection, and downstream analytical framework. A total of 510 studies were identified through structured searches for epilepsy- and SUDEP-related genomic data. After applying exclusion criteria (duplicate records, insufficient genomic annotation, non-exonic variants, unclear phenotypes, and lack of original sequencing data), 419 studies were retained, comprising 34,421 epilepsy cases (2,753 genes) and 1,238 SUDEP cases (896 genes). Following variant annotation and filtering, 1,630 epilepsy-associated and 544 SUDEP-associated genes were retained, yielding 33 overlapping genes with recurrent variants. Subsequent analyses included variant characterization and integrative multi-omic profiling.

### Demographic and Clinical Landscape of the SUDEP Cohort

Demographic information was incompletely reported across the SUDEP literature. Where available, sex was reported for 155 cases, revealing a male predominance: 91 males (59%) and 64 females (41%) (Supplementary Figure S1a and S1b). Age was reported for 236 cases, with infants (27%) and children (25%) comprising the largest groups. The remainder of the cohort included adults aged 30–49 years (20%), young adults 18–29 years (14%), adults 50–59 years (7%), adolescents 12–17 years (4%), and individuals aged ≥60 years (3%) (Supplementary Figure S1c). Given the extent of missing metadata, these distributions should be interpreted cautiously.

Comorbidity data were reported for only 187 of 1,238 SUDEP cases. Among those, neurological and cardiovascular conditions were most frequently documented (Supplementary Figure S1d). Neurological comorbidities were present in 160/187 cases, most commonly epilepsy (56.3%; 90/160) and seizure diagnoses (26.9%; 43/160). Less frequently reported neurological conditions included developmental delay and language impairment (8.8%), Dravet syndrome (6.9%), West syndrome (0.6%), and Ohtahara syndrome (0.6%). Cardiovascular comorbidities were noted in 14% (27/187) of cases and spanned a range of diagnoses, including Long QT syndrome (33.3%), arrhythmogenic right ventricular cardiomyopathy (18.5%), dilated and hypertrophic cardiomyopathies (14.8%), myocarditis (11.1%), Brugada syndrome (7.4%), venous malformations (7.4%), sick sinus syndrome (3.7%), and congenital cardiac anomalies (3.7%).

### Variant Classification and Mutation Spectrum in the SUDEP and Epilepsy Cohorts

Variant annotation was conducted in accordance with ACMG guidelines (Richards et al., 2015), focusing on rare variants with a gnomAD allele frequency below 0.001, CADD scores of 20 or higher, and excluding those classified as benign or likely benign in ClinVar. Following this filtering, the dataset was refined to include 698 variants across 544 genes in the SUDEP cohort and 21,943 variants across 1,630 genes in the epilepsy cohort. Within the SUDEP dataset, 92.1% (643/698) of variants were classified as VUS, while the remaining 7.9% were pathogenic or likely pathogenic (Figure 2a). An assessment of genomic localization of these rare clinically relevant variants demonstrated a predominant localization in exonic regions in exonic regions across both cohorts, with 97.4% of variants in SUDEP (680/698) and 96.9% (21,273/21,943) in epilepsy cases. In contrast, only a small fraction localized to splice sites, representing 2.5% in SUDEP and 3.0% in epilepsy (Figure 2b & 2c). Fisher’s exact test indicated that the differences in variant distribution between exonic and splice-site regions were not statistically significant between the cohorts (OR=1.18; 95% CI: 0.74–1.91; p=0.574).

**Figure 2.**
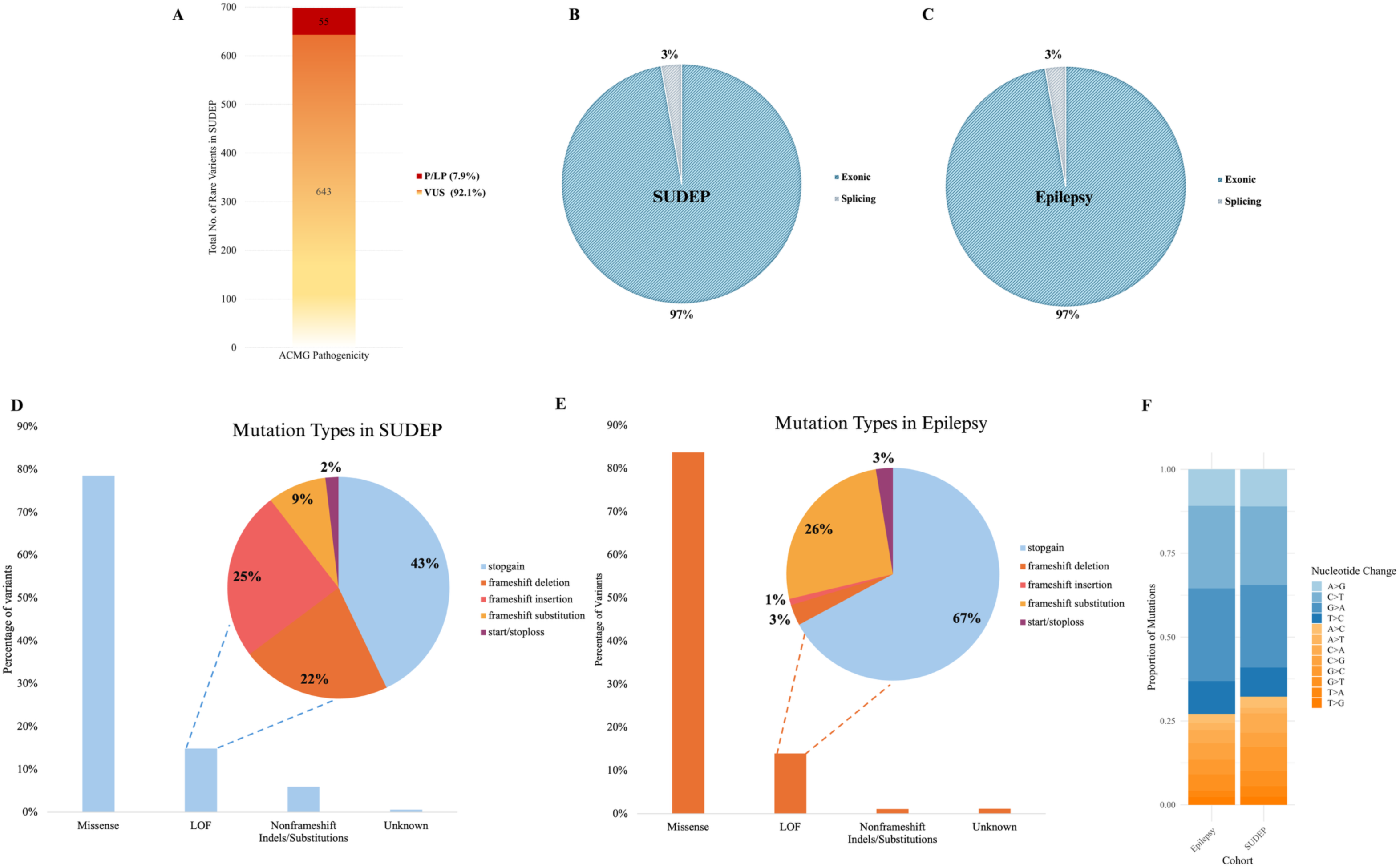
Variant classification and mutational spectrum in SUDEP and epilepsy cohorts. (a) ACMG-based pathogenicity classification showing predominance of variants of uncertain significance (VUS; 92.1%) in SUDEP, with 7.9% classified as pathogenic or likely pathogenic (P/LP). (b–c) Genomic localization of rare variants demonstrating enrichment in exonic regions in both SUDEP (97.4%) and epilepsy (96.9%) cohorts, with a small fraction in splice-site regions (∼3%). (d–e) Distribution of variant types across cohorts. (f) Mutational spectrum showing comparable nucleotide substitution profiles between SUDEP and epilepsy.

Further evaluation of variant classes within the exonic variant category revealed that missense variants were the predominant type in both groups, accounting for 78.4% (533/680) in SUDEP and 84.4% (17,963/21,273) in epilepsy (Figure 2d & 2e). The remaining variants were largely loss-of-function (LOF), comprising 15.4% (105/680) in SUDEP and 13.3% (2,836/21,273) in epilepsy, although these differences were not statistically significant (OR=1.19; 95% CI: 0.96–1.47; p=0.12). Non-frameshift indels and substitutions appeared more prevalent in SUDEP (5.6%, 38/680) compared with epilepsy (0.9%, 195/21,273), whereas approximately 1% variants in each cohort were categorized as unknown. Within the LOF subset, stop-gain variants constituted the largest proportion in both cohorts (OR=0.47; 95% CI: 0.32–0.70; p=2.0 × 10⁻⁴), representing 48% (48/105) of SUDEP LOF variants and 66% (1,634/2,836) of epilepsy LOF variants (Figure 2d & 2e). This pattern indicates a higher relative proportion of frameshift variants in SUDEP, whereas stop-gained variants show a higher occurrence in epilepsy.

The mutational spectrum analysis demonstrated that both cohorts were dominated by C>T transitions, followed by G>A and T>C substitutions. Transversion events were less frequent and showed broadly comparable distributions between cohorts, with only minor differences observed across specific nucleotide changes (Figure 2f).

### Shared Genetic Signatures of Epilepsy and SUDEP

Out of 1,630 epilepsy-associated genes and 544 SUDEP-associated genes, we first sought to identify the overlapping set that may contribute to the shared biology between the two phenotypes. Among 149 shared genes between the two conditions, 33 carried two or more rare variants in both epilepsy and SUDEP unrelated cases (Figure 3a). These recurrently mutated genes spanned several functional categories, including ion channels and transporters such as *SCN1A*, *SCN2A*, *SCN5A*, *SCN10A*, *KCNE1*, *KCNH2*, and *KCNA1*; pacemaker and calcium regulators such as *HCN1*, *HCN2*, *HCN4*, *CACNA1A*, *CACNA1H*, and *RYR2*; structural and contractile proteins including *TTN*, *MYH6*, and *MYO18B*; metabolic and lysosomal enzymes such as *DHTKD1*, *ALDH5A1*, *SCARB2*, and *PIGA*; synaptic and signaling mediators such as *GRIN2B*, *PRRT2*, and *ADGRV1*; and regulatory or repair-associated genes including *DEPDC5* and *PNKP*. Notably, variants in ion channel and transporter genes accounted for the largest proportion of variants across these 33 overlapping genes.

**Figure 3.**
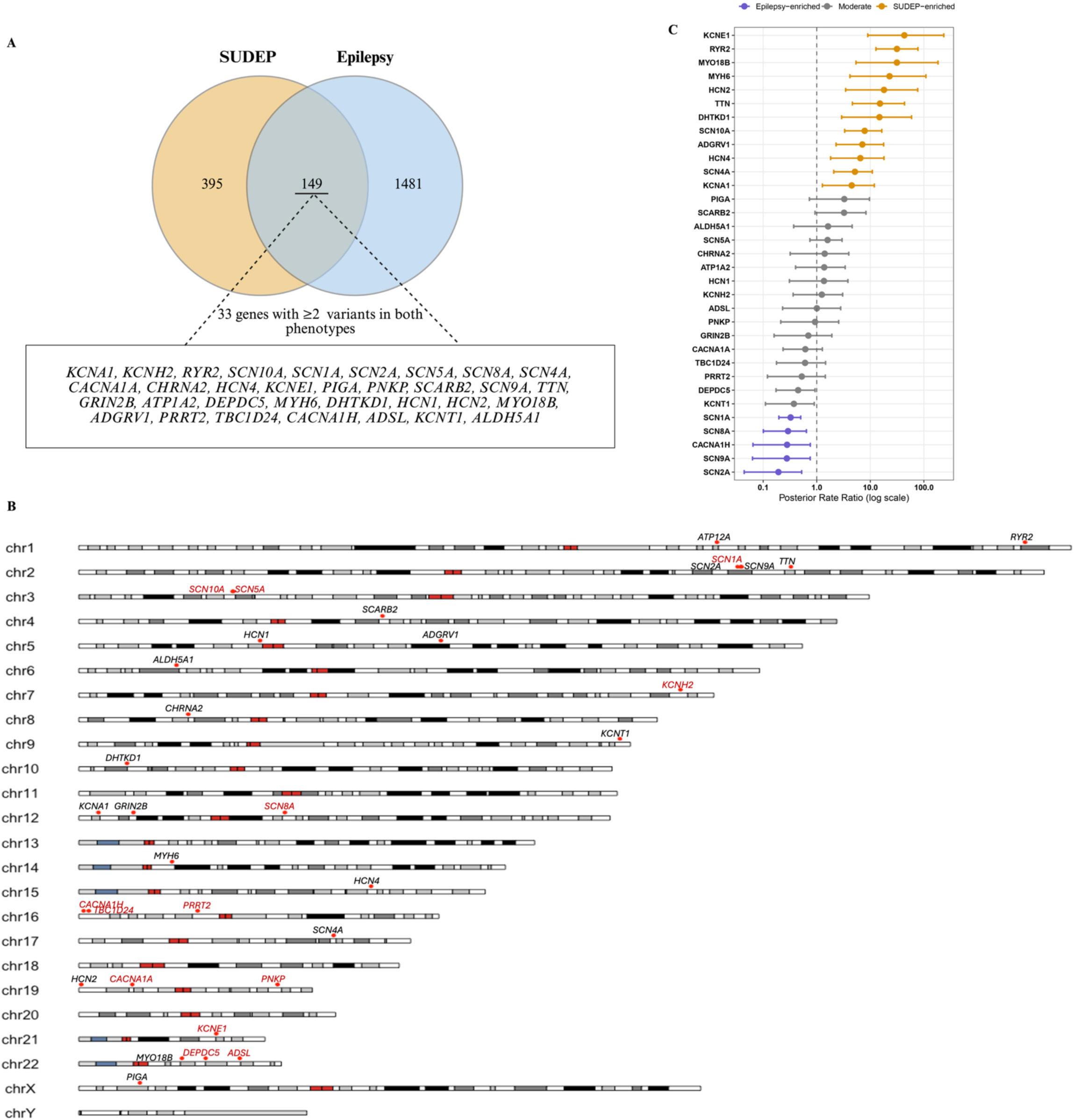
Shared and distinct genetic architecture of epilepsy and SUDEP. (a) Venn diagram showing the overlap between epilepsy- and SUDEP-associated genes. Out of 1,630 epilepsy and 544 SUDEP genes, 149 were shared, with 33 genes carrying two or more variants in both phenotypes. (b) Chromosomal distribution of the 33 overlapping genes, highlighting recurrent loci on chromosomes 2 (*SCN1A, SCN2A, SCN9A, TTN*), 3 (*SCN5A, SCN10A*), and 12 (*SCN8A, GRIN2B, KCNA1*). Genes highlighted in red indicate those harboring the same recurrent variants in both cohorts. (c) Bayesian rate ratio analysis depicting gene-level enrichment between cohorts. Genes such as *KCNE1, RYR2, MYO18B, MYH6, HCN2, TTN*, *DHTKD1, SCN10A, ADGRV1, HCN4, SCN4A,* and *KCNA1* were significantly SUDEP-enriched (yellow), while *SCN1A, SCN8A, CACNA1H, SCN9A* and *SCN2A* were epilepsy-enriched (purple). Grey points denote genes with non-significant differences (95% credible intervals spanning 1). Error bars represent 95% credible intervals.

Chromosomal mapping showed a wide genomic distribution, with recurrent loci on chromosome 2 (*SCN1A*, *SCN2A*, *SCN9A*, *TTN*), chromosome 3 (*SCN5A*, *SCN10A*), and chromosome 12 (*SCN8A*, *GRIN2B*, *KCNA1*) (Figure 3b). Several recurrent variants were also detected across unrelated SUDEP and epilepsy cases, primarily affecting genes contributing to ion channel activity and excitability regulation (Table 1). Comparative analysis also revealed that *SCN1A*, *SCN5A*, *SCN8A*, *KCNE1*, and *RYR2* harbored several pathogenic or likely pathogenic variants shared between epilepsy and SUDEP, whereas additional variants in *PRRT2*, *PNKP*, and *ADSL* linked metabolic and synaptic pathways across the two phenotypes.

**Table 1.** Shared pathogenic variants reported in SUDEP and epilepsy. Recurrently reported variants identified in SUDEP and/or epilepsy cases, including their genomic annotation, predicted pathogenicity, and reporting studies, highlighting overlapping loci implicated in neuronal excitability and cardiac conduction.

| Gene | Variant Effect | NM_Accession | Codon Change | A.A Change | CLNSIG | CADD | REVEL | gnomAD_AF | Annotation | Reference paper (Epilepsy/SUDEP) |
| --- | --- | --- | --- | --- | --- | --- | --- | --- | --- | --- |
| *ADSL* | Missense | NM_000026 | c.G1288A | p.D430N | VUS | 26.3 | 0.659 | 1.32E-05 | VUS | (Palmer et al., 2018)/(Coll et al., 2017) |
| *DEPDC5* | Stopgain | NM_001369901 | c.C646T | p.Q216X | P | 35 | . | 0.00 | P/LP | (Heyne et al., 2019)/(Nascimento et al., 2015) |
| *CACNA1A* | Missense | NM_001127221 | c.C2407A | p.R803S | CI | 24.3 | 0.496 | 1.31E-05 | VUS | (Truty et al., 2019)/(Aschner et al., 2024) |
|  |  |  | c.C6724T | p.R2242W | VUS | 22.9 | 0.467 | 6.58E-06 | VUS | (Truty et al., 2019)/(Aschner et al., 2024) |
| *CACNA1H* | Missense | NM_001005407 | c.C4709T | p.A1570V | VUS | 33 | 0.659 | 4.60E-05 | VUS | (Truty et al., 2019)/(Coll et al., 2017) |
|  |  |  | c.G5006A | p.R1669Q | VUS | 32 | 0.804 | 8.54E-05 | VUS | (Tozkir et al., 2025; Truty et al., 2019)/(Coll et al., 2016) |
| *KCNE1* | Missense | NM_001127670 | c.G253A | p.D85N | CI | 21.7 | 0.33 | 0.00 | VUS | (Partemi et al., 2015)/(Hata et al., 2017) |
| *KCNH2* | Missense | NM_000238 | c.T245C | p.I82T | LP | 24.5 | 0.787 | 0.00 | P/LP | (Partemi et al., 2013)/(Partemi et al., 2015) |
| *SCN1A* | Missense | NM_001353948 | c.A3924T | p.E1308D | CI | 22.8 | 0.499 | 6.00E-04 | P/LP | (Till et al., 2020)/(Rochtus et al., 2020) |
|  |  |  | c.T5422C | p.F1808L | P | 27.3 | 0.976 | 0.00 | P/LP | (Fujiwara et al., 2003; Ishii et al., 2017)/(Aschner et al., 2024) |
|  |  |  | c.G235C | p.D79H | . | 25.6 | 0.956 | 0.00 | P/LP | (Harkin et al., 2007)/(Cooper et al., 2016) |
|  | Stopgain |  | c.C1837T | p.R613X | P | 35 | . | 0.00 | P/LP | (Ishii et al., 2017; Negi et al., 2024; Truty et al., 2019)/(Cooper et al., 2016; Leu et al., 2015) |
| *SCN5A* | Missense | NM_001099405 | c.C5440G | p.Q1814E | CI | 23 | 0.53 | 4.00E-04 | VUS | (Partemi et al., 2015)/(Coll et al., 2016) |
|  | Missense | NM_001160161 | c.C3415T | p.R1139W | VUS | 22.9 | 0.647 | 5.25E-05 | VUS | (Truty et al., 2019)/(Ge et al., 2020) |
| *SCN8A* | Missense | NM_001177984 | c.C2300T | p.T767I | P/LP | 25.8 | 0.967 | 0.00 | P/LP | (Heyne et al., 2019; Lindy et al., 2018) /(Fang et al., 2023) |
|  |  |  | c.A5179G | p.N1727D | P | 25.7 | 0.912 | 0.00 | P/LP | (Zhu et al., 2017)/(Veeramah et al., 2012) |
| *SCN10A* | Missense | NM_001293307 | c.T3506G | p.M1169R | VUS | 26.6 | 0.965 | 1.31E-05 | VUS | (Fernández-Marmiesse et al., 2019)/(Coll et al., 2016) |
| *SCARB2* | Missense | NM_001204255 | c.A194G | p.Y65C | VUS | 29.9 | 0.876 | 2.00E-04 | VUS | (Truty et al., 2019)/(Aschner et al., 2024) |
|  |  |  | c.G277A | p.E93K | VUS | 32 | 0.778 | 2.00E-04 | LP | (Truty et al., 2019)/(Coll et al., 2017) |
| *PRRT2* | Stopgain | NM_001256442 | c.C718T | p.R240X | P | 35 | . | 0.00 | P/LP | (Heyne et al., 2019; Labate et al., 2013)/(Labate et al., 2013) |
| *PNKP* | Missense | NM_007254 | c.T1397C | p.M466T | VUS | 22.4 | 0.129 | 6.03E-05 | VUS | (Truty et al., 2019)/(Aschner et al., 2024) |
| *TBC1D24* | Missense | NM_001199107 | c.G328A | p.G110S | CI | 25 | 0.745 | 0.00 | VUS | (Balestrini et al., 2016; Truty et al., 2019)/(Trivisano et al., 2017) |

To statistically compare the phenotype-specific burden associated with each shared gene, we applied a Bayesian Poisson–Gamma model to estimate gene-specific posterior rate ratios (RR) (Figure 3c, Supplementary Table 3). This probabilistic framework allowed direct comparison of variant burden per case between SUDEP and epilepsy while accounting for differences in cohort size, sparse counts, and uncertainty inherent to rare variants. Genes meeting the criteria for SUDEP-specific enrichment (posterior RR > 1, 95% credible interval not spanning 1, and FDR < 0.05) included *KCNE1*, *RYR2*, and *TTN* as key cardiac effectors (all FDR ≤ 1.65 × 10⁻⁴). Additional SUDEP-enriched genes included *MYO18B*, *MYH6*, *HCN2*, *DHTKD1*, *SCN10A*, *ADGRV1*, *HCN4*, *SCN4A*, and *KCNA1*, supporting the involvement of cardiac excitability, pacemaker function, and contractile pathways in SUDEP risk. In contrast, canonical neuronal ion channel genes such as *SCN1A, SCN2A, SCN8A, SCN9A,* and *CACNA1H* showed enrichment in epilepsy (RR < 1 with credible intervals not spanning 1), while several established epilepsy-associated genes, including *CACNA1A, KCNT1, GRIN2B, PRRT2, DEPDC5,* and *TBC1D24* did not show significant differential enrichment between cohorts, indicating similar variant burden between SUDEP and epilepsy.

### Domain-Level Variant Mapping Identifies Shared and SUDEP-Specific Hotspots

The mutational burden analysis across epilepsy and SUDEP cohorts revealed distinct variant patterns, reinforcing channelopathies as a major determinant of risk (Supplementary Figure 2). Domain-level evaluation of the 33 overlapping genes showed both shared and phenotype-associated substitutions in *ADGRV1*, *ADSL*, *ATP1A2*, *ALDH5A1*, and *DEPDC5*. Notably, *DEPDC5* exhibited recurrent clustering of variants at specific positions, along with variants observed in SUDEP cases, consistent with its established role in mTOR pathway–mediated epileptogenesis (De Fusco et al., 2020).

As described earlier, ion channel genes carried the largest burden of rare variants. *CACNA1A*, *CACNA1H*, and *CHRNA2* displayed dense clustering of substitutions across annotated functional domains, with variants observed in both epilepsy and SUDEP cases. Across sodium and potassium channel families, recurrent substitutions were detected in *KCNE1*, *KCNA1*, *KCNH2*, *KCNT1*, *SCN1A*, *SCN2A*, *SCN5A*, *SCN8A*, and *SCN10A*, frequently mapping to annotated functional regions. In particular, *KCNT1* and *SCN1A* showed high variant density, including both shared and phenotype-associated clustering patterns.

Beyond classical channel genes, several neuro-cardiac regulators also demonstrated convergent patterns. *GRIN2B* variants clustered within annotated functional domains, including regions corresponding to ligand-binding and transmembrane segments, with additional variants observed in SUDEP cases. Similarly, variants in *HCN1*, *HCN2*, and *HCN4* localized to pore and cyclic nucleotide-binding regions, consistent with potential effects on pacemaker channel function. Variants in *KCNE1* and *KCNH2* were observed within regions critical for channel gating and repolarization, aligning with their known roles in cardiac electrophysiology and arrhythmia susceptibility.

Outside ion/transport channels, recurrent substitutions were also identified in structural and synaptic genes. *RYR2* variants spanned domains critical for calcium release and excitation–contraction coupling. *MYO18B*, *MYH6*, and *TTN* carried rare substitutions altering sarcomeric function, while *PRRT2* showed a pronounced hotspot at c.640delinsGC, the most frequent variant observed in the dataset. *PNKP* variants clustered within functionally annotated regions consistent with its role in DNA repair, whereas *PIGA* and *TBC1D24* harbored recurrent mutations within domains essential for GPI-anchor biosynthesis and synaptic vesicle trafficking, respectively.

These observations indicate that while ion channels represent the predominant carriers of rare deleterious variants, recurrent substitutions in genes regulating calcium handling, contractile stability, synaptic function, and DNA repair broaden the mechanistic landscape underlying epilepsy and SUDEP-associated risk.

### Functional Enrichment and Pathway Architecture of Shared and SUDEP-Specific Genes

Functional enrichment analysis of the shared gene set identified four major interconnected clusters: Ion Channel Activity, Neuro–Cardiac Response & Regulation, Signaling Regulation, and Contractile Dynamics (Figure 4a). Ion Channel Activity showed the strongest enrichment and included pathways related to voltage-gated ion channel activity, membrane depolarization, regulation of membrane potential, and action potential propagation. These pathways were driven largely by genes such as *SCN1A, SCN2A, SCN5A, SCN8A, CACNA1A*, and *KCNE1*, underscoring the central role of channelopathies in both epilepsy and SUDEP (Bagnall et al., 2016; Sahly et al., 2022).

**Figure 4.**
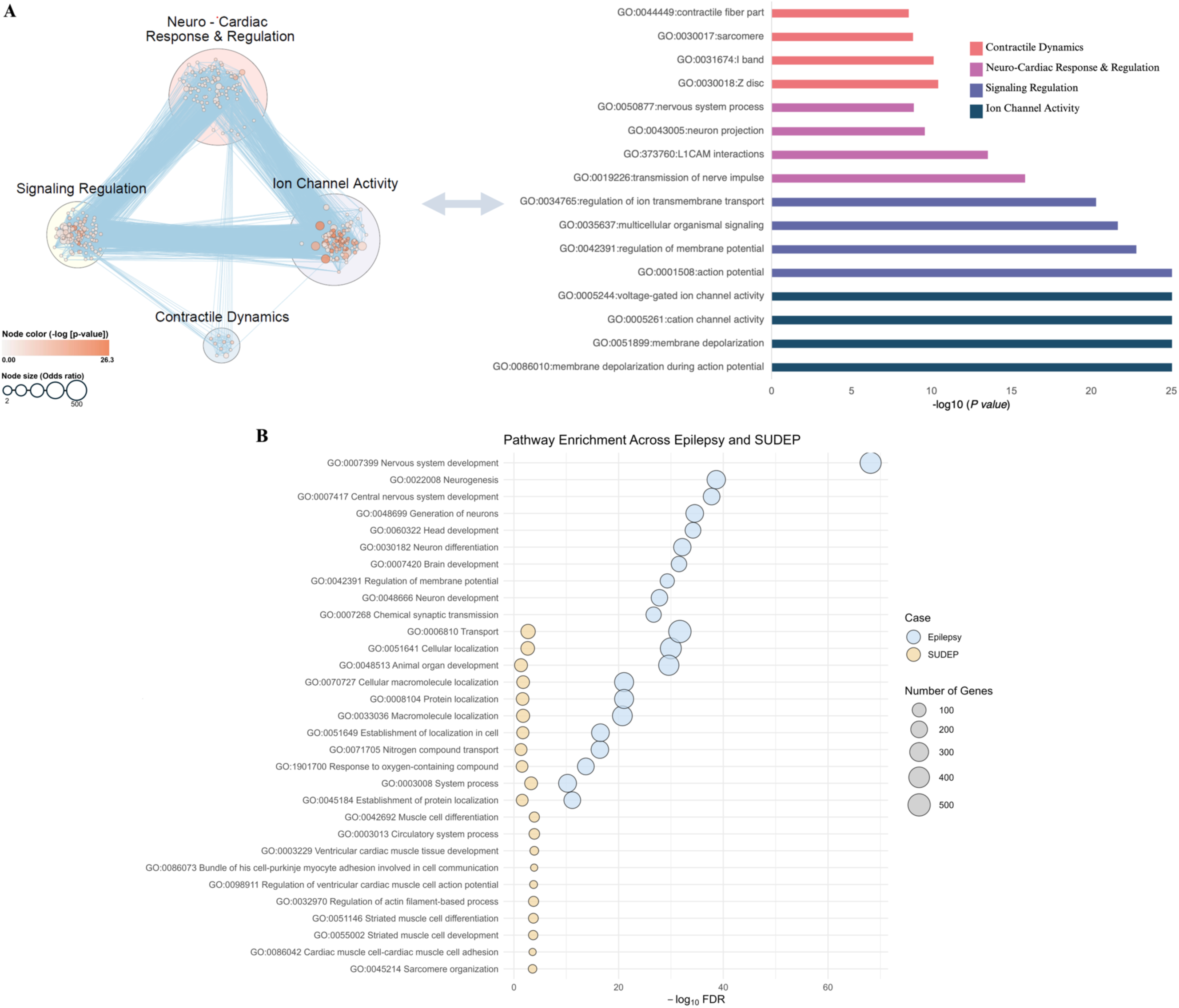
Functional enrichment and pathway architecture of shared and distinct genes in epilepsy and SUDEP. (a) Network-based Gene Ontology (GO) enrichment analysis of shared genes revealed four interconnected clusters: Ion Channel Activity, Neuro–Cardiac Response & Regulation, Signaling Regulation, and Contractile Dynamics. The most significantly enriched pathways were voltage-gated ion channel activity, membrane depolarization, and regulation of membrane potential, followed by neuronal projection and sarcomere organization, highlighting the convergence of excitability and contractile processes. Node size reflects odds ratio, and node color intensity corresponds to statistical significance (–log *p*-value). (b) Comparative pathway enrichment between epilepsy (blue) and SUDEP (yellow) cohorts. Epilepsy-specific pathways were enriched for neurodevelopmental and synaptic signaling processes whereas SUDEP-specific pathways showed strong enrichment for cardiac conduction and muscle structure organization. Shared pathways included regulation of membrane potential and ion transport, reflecting overlapping neuro-cardiac excitability mechanisms.

The Neuro–Cardiac Response & Regulation cluster consisted of pathways associated with nerve impulse transmission, neuron projection, L1CAM-mediated cell adhesion, and multicellular signaling. These pathways highlight shared neuro-cardiac regulatory processes influencing both cortical excitability and cardiac rhythm stability. The Contractile Dynamics module included sarcomere organization, contractile fiber components, and Z-disc assembly, suggesting a potential contribution of myocardial structural organization and excitation–contraction coupling to SUDEP risk. Meanwhile, the Signaling Regulation cluster captured pathways related to ion transport and intercellular communication, linking neuronal signaling mechanisms with cardiac electrophysiology.

Analysis of genes enriched in the SUDEP cohort revealed strong enrichment for cardiac conduction, sarcomeric, and muscle-specific processes (Figure 4b). Prominent pathways included bundle of His cell–Purkinje myocyte communication, regulation of cardiac muscle action potential, striated muscle cell differentiation, sarcomere organization, and actin filament–based processes. These findings point to a cardiac-focused molecular network influencing impulse propagation and contractile function, consistent with arrhythmogenic mechanisms reported in SUDEP (Muccioli et al., 2024; Ryvlin et al., 2013; Van der Lende et al., 2016) and excitation–contraction defects in SUDEP (Ryvlin et al., 2013; Sahly et al., 2022). In contrast, genes enriched in epilepsy were associated with neurodevelopmental and synaptic signaling pathways (Boulaki et al., 2025), including nervous system development, neurogenesis, chemical synaptic transmission, and regulation of membrane potential (Figure 4b). These pathways reinforce the dominant role of neurodevelopmental and neurotransmission-related processes in epileptogenesis and seizure susceptibility.

Importantly, several pathways, such as regulation of membrane potential, nerve impulse transmission, and multicellular signaling, were shared between epilepsy and SUDEP, indicating mechanistic overlap centered on ion flux regulation and neuronal–cardiac excitability. However, enrichment in the SUDEP cohort extended beyond shared excitability pathways to include myocardial conduction, contractile fiber organization, and sarcomeric structure, highlighting a distinct cardiac component within the broader shared channelopathy landscape.

### Tissue and Isoform-Level Expression Landscapes of Shared Epilepsy–SUDEP Genes Across Brain and Heart

To characterize how genes shared between epilepsy and SUDEP are deployed across organs involved in neuro–cardiac regulation, we examined their tissue-level transcriptional profiles using GTEx RNA-seq datasets (https://gtexportal.org/) from human brain and heart (Figure 5, Supplementary Figure 3). This analysis revealed clear tissue-specific biases that distinguish neuronal and cardiac functional programs within the shared genetic landscape of epilepsy and SUDEP. Cardiac excitability and contractile genes, including *MYH6, TTN, MYO18B, RYR2,* and *SCN5A*, showed strong heart-enriched expression, whereas neuronal ion channel and synaptic signaling genes such as *SCN1A, SCN2A, HCN2, GRIN2B*, and *PRRT2* were strongly brain-enriched. Notably, several genes exhibited relatively balanced expression across both tissues, indicating shared functional roles across the neuro–cardiac axis.

**Figure 5.**
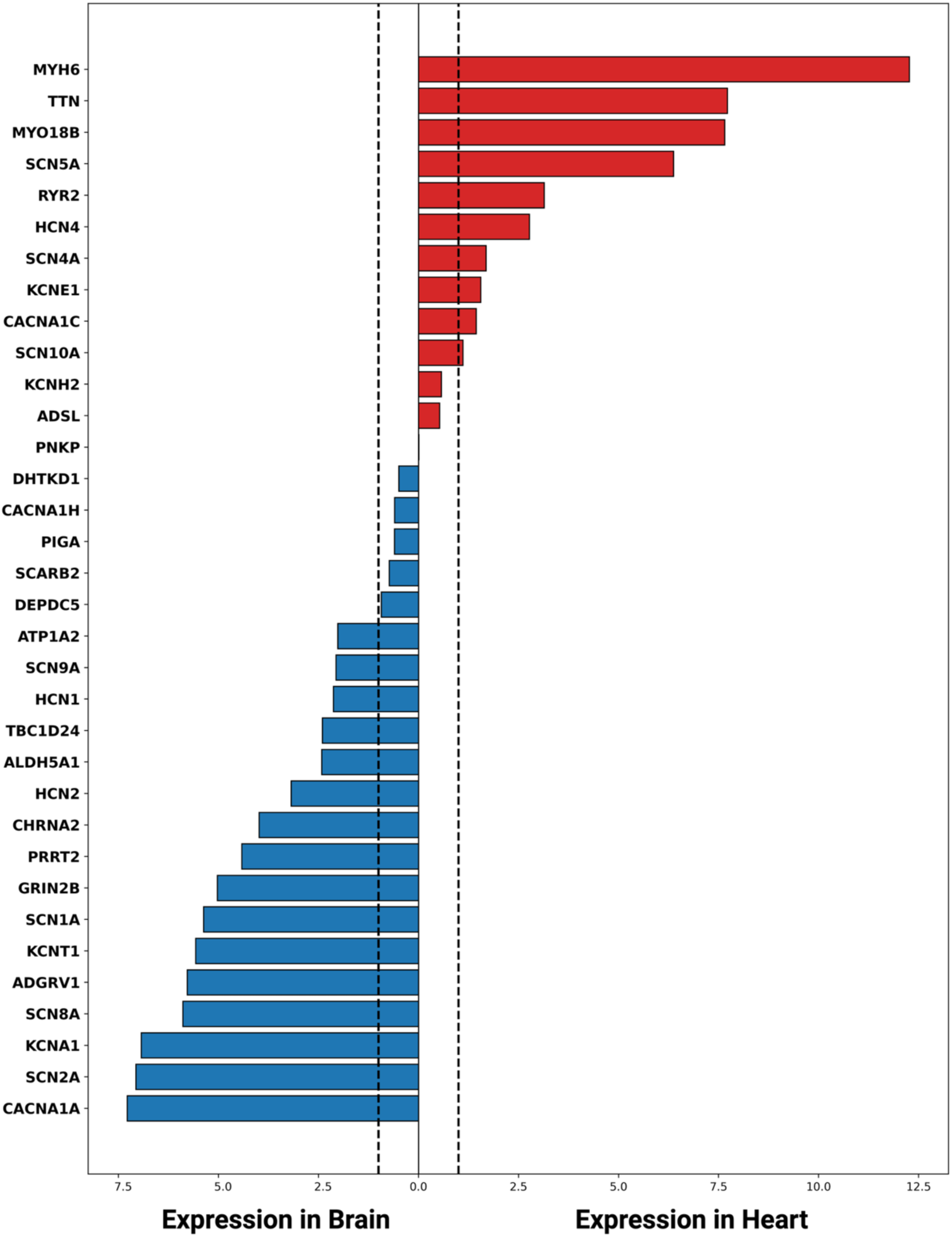
Tissue-specific expression patterns of shared epilepsy–SUDEP genes. Comparative expression analysis of overlapping genes across human brain and heart tissues from GTEx RNA-seq data. Positive log₂ fold-change (log₂FC) values denote heart-enriched expression, observed prominently for *MYH6, TTN, MYO18B, RYR2*, and *SCN5A*, whereas negative values represent brain-enriched expression seen in neuronal and synaptic genes such as *SCN1A, SCN2A, HCN2*, and *GRIN2B*. Vertical dashed lines indicate enrichment thresholds (log₂FC = ±1), corresponding to at least a twofold difference in expression between tissues.

To further resolve how these genes are regulated within individual cell populations and at the level of transcript isoforms, we next analyzed publicly available single-cell long-read isoform atlases from human heart and brain tissues (Hardwick et al., 2022; Pan et al., 2025). Following preprocessing and integration of the datasets, cellular populations were defined using canonical lineage markers to identify the major cell types present in each tissue (Figure 6). In the heart dataset, the principal populations included cardiomyocytes, endothelial cells, fibroblasts, pericytes, macrophages, T cells, smooth muscle cells, and endocardial and lymphatic endothelial compartments. In the brain dataset, cell types comprised excitatory neurons, inhibitory neurons, astrocytes, oligodendrocytes, oligodendrocyte precursor cells (OPCs), microglia, and VIM⁺ cell populations.

**Figure 6.**
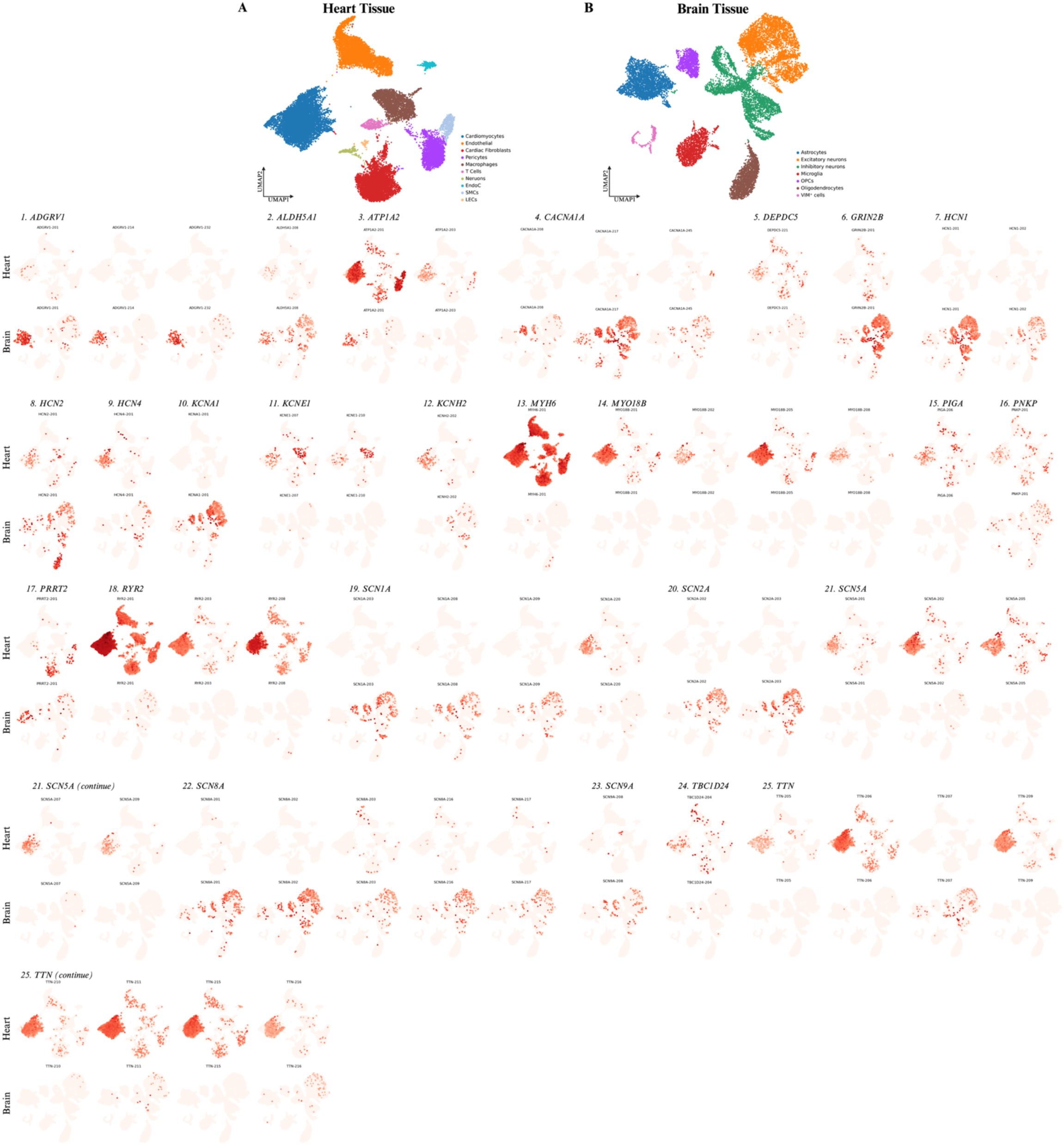
Tissue- and cell-type–resolved expression patterns of epilepsy–SUDEP–associated genes across human heart and brain. (A) UMAP representation of single-cell transcriptomic profiles from human heart tissue, showing major cardiac cell populations including cardiomyocytes, endothelial cells, fibroblasts, pericytes, smooth muscle cells, and immune cell populations. (B) UMAP representation of brain tissue single-cell transcriptomes highlighting major neuronal and glial populations, including excitatory neurons, inhibitory neurons, astrocytes, oligodendrocytes, microglia, and OPC cells. Lower panels display gene expression projections for 25 genes associated with epilepsy and SUDEP risk across the two tissues. For each gene, expression intensity is shown as red signal overlaid on the UMAP embeddings, with separate rows for heart and brain datasets to illustrate tissue-specific distribution. The analyzed genes include *ADGRV1, ALDH5A1, ATP1A2, CACNA1A, DEPDC5, GRIN2B, HCN1, HCN2, HCN4, KCNA1, KCNE1, KCNH2, MYH6, MYO18B, PIGA, PNKP, PRRT2, RYR2, SCN1A, SCN2A, SCN5A, SCN8A, SCN9A, TBC1D24,* and *TTN*. These maps reveal pronounced tissue- and cell-type–specific expression patterns, with neuronal ion channel genes, for instance *ADGRV1, CACNA1A, HCN1, GRIN2B, SCN1A, SCN2A, SCN8A, SCN9A* enriched in neuronal populations in brain tissue, while structural and electrophysiological genes associated with cardiac function, including *ATP1A2, MYH6, MYO18B, RYR2*, *SCN5A, TTN* show strong enrichment in cardiac tissues.

Using these annotated cell populations, we examined isoform-resolved expression patterns across candidate genes associated with SUDEP. For each gene, individual transcript isoforms were analyzed independently to determine whether expression patterns were shared across tissues, restricted to a single tissue, or partitioned between tissues through alternative transcript expression. To ensure robustness and minimize low-frequency transcriptional noise, transcripts were considered expressed only if they were detected in at least 100 cells, annotated as protein-coding based on Ensembl biotype classification, and supported by more than 10 UMI counts. Importantly, lack of detected expression should not be interpreted as true absence, but may reflect technical limitations of single-cell sequencing, including dropout events and limited detection sensitivity.

Across datasets, genes segregated into three principal patterns of transcript expression: (i) heart-restricted isoforms, (ii) brain-restricted isoforms, and (iii) comparative isoform expression across tissues (Figure 6, Table 2). These patterns indicate that gene sharing between epilepsy and SUDEP occurs at the locus level, whereas functional deployment is frequently partitioned at the level of transcript isoforms.

**Table 2.** Tissue-specific isoform architecture of epilepsy–SUDEP genes across human heart and brain. Isoform-resolved expression patterns of candidate epilepsy–SUDEP genes derived from publicly available long-read single-cell transcriptomic atlases of human heart and brain tissues. Only protein-coding transcripts (Ensembl v110) were considered. A transcript was classified as expressed if detected in ≥100 cells and supported by ≥10 UMI counts. Genes were categorized into four expression patterns: heart-restricted (isoforms detected exclusively in cardiac cell populations), brain-restricted (isoforms detected only in neuronal or glial populations), comparative isoforms (distinct isoforms detected in heart and brain), and comparable isoforms (shared transcript isoforms detected across both tissues). Absence of detection does not imply true absence but may reflect technical limitations of single-cell sequencing.

| Gene | Heart Isoforms | Brain Isoforms | Shared Isoforms |
| --- | --- | --- | --- |
| Heart-restricted |  |  |  |
| *HCN4* | 201 | – | – |
| *KCNH2* | 202 | – | – |
| *KCNE1* | 207, 210 | – | – |
| *MYH6* | 201 | – | – |
| *MYO18B* | 201, 202, 205, 208 | – | – |
| *PIGA* | 206 | – | – |
| *PNKP* | 201 | – | – |
| *PRRT2* | 201 | – | – |
| *ATP1A2* | 201, 203 | – | – |
| Brain-restricted |  |  |  |
| *ADGRV1* | – | 201, 214, 232 | – |
| *CACNA1A* | – | 208, 217, 245 | – |
| *GRIN2B* | – | 201 | – |
| *HCN1* | – | 201, 202 | – |
| *HCN2* | – | 201 | – |
| *KCNA1* | – | 201 | – |
| *SCN1A* | – | 203, 208, 209, 216 | – |
| *SCN2A* | – | 202, 203 | – |
| *SCN8A* | – | 201, 202, 203, 216, 217 | – |
| Comparative isoforms |  |  |  |
| *ALDH5A1* | – | 208 | 202 |
| *DEPDC5* | 221 | – | 208 |
| *RYR2* | 201, 203, 208 | – | 207 |
| *SCARB2* | – | 227 | 201 |
| *SCN5A* | 201, 202, 205, 207, 209 | – | 204 |
| *SCN9A* | 211 | 208 | 202 |
| *TBC1D24* | 204 | – | 208 |
| *TTN* | 205, 206, 209, 210, 211,<br>215, 216 | 207 | 202, 204, 212, 214 |

#### Heart-Restricted Isoform Programs Reflect Cardiac Electrophysiology and Contractile Function

A subset of genes demonstrated isoforms detected exclusively in cardiac cell populations under the applied thresholds, consistent with roles in cardiac electrophysiology and contractile machinery. These included *HCN4, KCNH2, KCNE1, MYH6, MYO18B, ATP1A2, PIGA, PNKP*, and *PRRT2*.

Among these, HCN4-201 was confined to cardiac cells, aligning with its role in generating the pacemaker current (If) in the sinoatrial node (Fenske et al., 2020; Li et al., 2015).. Similarly, KCNH2-202 and KCNE1 isoforms (KCNE1-207 and -210), key components of cardiac repolarization, were restricted to heart datasets (Jonsson et al., 2012; Wang et al., 2020). Structural genes such as MYH6-201 and multiple *MYO18B* isoforms (MYO18B-201, -202, - 205, and -208) were likewise detected exclusively in cardiomyocytes, consistent with sarcomeric specialization (Granados-Riveron et al., 2010; Latham et al., 2020).

Notably, several genes with established roles outside the heart, including *ATP1A2* (Isaksen & Lykke-Hartmann, 2016; Rindler et al., 2013) and *PRRT2* (Döring et al., 2020; Huang et al., 2021), exhibited cardiac-restricted isoforms in this dataset, indicating context-dependent transcript deployment rather than canonical tissue assignment. These findings suggest that isoform-level regulation refines functional specialization within cardiac cell populations.

#### Brain-Restricted Isoforms Define Neuronal Excitability and Synaptic Programs

In contrast, a distinct set of genes exhibited isoforms restricted to neuronal and glial populations. These included *ADGRV1, CACNA1A, GRIN2B, HCN1, HCN2, KCNA1, SCN1A, SCN2A*, and *SCN8A*.

*ADGRV1* displayed multiple neuronal isoforms (ADGRV1-201, -214, and -232), reflecting transcript diversity in synaptic adhesion and signaling pathways (Lee et al., 2025). Ion channel genes central to neuronal excitability showed strong isoform restriction, including *HCN1* (HCN1-201, -202), *HCN2* (HCN2-201), and *KCNA1* (KCNA1-201) (Tang et al., 2008; Shah & Aizenman, 2014). Similarly, sodium channel genes demonstrated pronounced neuronal isoform specificity, with *SCN1A* (SCN1A-203, -208, -209, and -216), *SCN2A* (SCN2A-202 and -203), and *SCN8A* (SCN8A-201, -202, -203, -216, and -217) detected exclusively in brain datasets (Escayg & Goldin, 2010; Krause et al., 2015; Meisler & Kearney, 2005).

Calcium channel gene *CACNA1A* exhibited multiple neuronal isoforms (CACNA1A-208, - 217, and -245), while GRIN2B-201 was restricted to neuronal populations, consistent with its role in excitatory synaptic transmission. These patterns indicate that neuronal excitability genes are deployed through highly specific transcript isoforms within brain cell populations (Szymanowicz et al., 2024; J. Betzenhauser et al., 2015; Sabo et al., 2023).

#### Comparative Isoform Expression Reveals Transcript Partitioning Across Tissues

A major feature of the dataset was the presence of genes expressed in both tissues but utilizing distinct transcript isoforms, indicating isoform-level partitioning across the neuro–cardiac axis.

This pattern was evident in ion channel genes. *SCN5A*, the principal cardiac sodium channel, expressed multiple cardiac isoforms (SCN5A-201, -202, -205, -207, and -209), while a distinct isoform (SCN5A-204) was detected across both tissues (Veerman et al., 2015). Similarly, *SCN9A* exhibited a brain-specific isoform (SCN9A-208), a cardiac isoform (SCN9A-211), and a shared transcript (SCN9A-202), demonstrating layered transcript expression within a single locus (Baker & Nassar, 2020; Barajas-Martinez et al., 2020).

Structural and contractile genes also showed marked isoform partitioning. *TTN* displayed extensive cardiac isoform diversity (TTN-205, -206, -209, -210, -211, -215, and -216) (Jolfayi et al., 2024), while TTN-207 was detected in neuronal populations (Cameron et al., 2024) and multiple isoforms (TTN-202, -204, -212, and -214) were shared between tissues. Similarly, *RYR2* expressed cardiac isoforms (RYR2-201, -203, and -208) alongside a shared isoform (RYR2-207), consistent with roles in both cardiac excitation–contraction coupling and neuronal calcium signaling (Yap & Smyth, 2019).

Additional genes, including *DEPDC5*, a regulator of the mTOR signaling pathway associated with epilepsy (Bacq et al., 2022; Groff et al., 2025) and the GABA metabolic gene *ALDH5A1* (Afshar-Saber et al., 2024), as well as *SCARB2* and *TBC1D24*, exhibited combinations of tissue-specific and shared isoforms. For instance, *SCARB2* showed a neuronal isoform (SCARB2-227) alongside a shared transcript (SCARB2-201), while *DEPDC5* displayed a cardiac-specific isoform (DEPDC5-221) in addition to a shared transcript (DEPDC5-208). These patterns indicate that isoform selection, rather than gene presence alone, governs tissue-specific functional deployment.

Integration of isoform-level expression patterns with gene-level enrichment analysis (Figure 3c) revealed concordant biological stratification. Genes enriched in epilepsy were predominantly brain-restricted or neuronally biased, including *SCN2A, SCN8A, KCNT1, GRIN2B, DEPDC5,* and *TBC1D24*, consistent with disruption of neuronal ion channel and synaptic signaling networks (Glasscock, 2014; Sun et al., 2023). In contrast, SUDEP-enriched genes showed convergence toward cardiac-expressed or cardiac-partitioned isoforms, including *KCNE1*, *HCN4, RYR2, TTN, MYH6*, and *MYO18B.* Many of these genes also exhibited heart-enriched or heart-restricted isoform expression, linking genetic enrichment to cardiac electrophysiological and contractile pathways. Importantly, comparative isoform expression was observed across both neuronal and cardiac gene sets, suggesting that disease divergence arises not only from gene-level enrichment but also from transcript-level regulation. Alternative isoform selection may alter channel kinetics, protein localization, or interaction networks without requiring changes in overall gene expression. Collectively, these findings demonstrate that shared genetic loci between epilepsy and SUDEP are differentially deployed across tissues through isoform-specific regulation. Epilepsy-associated risk is primarily associated with neuronal isoform programs governing excitability and synaptic transmission, whereas SUDEP-associated risk shows stronger alignment with cardiac isoforms regulating rhythm and contractile function (Mir et al., 2025; Arking & Sotoodehnia, 2012; Chahal et al., 2020; Shlobin et al., 2024). Rather than representing a more severe form of epilepsy, SUDEP emerges as a disorder of the neuro–cardiac interface, in which isoform-level partitioning functionally separates neuronal and cardiac programs within shared genes (Shlobin et al., 2024; Sun et al., 2023). This framework provides a mechanistic basis for how identical genetic loci can drive divergent phenotypic outcomes across organ systems.

### Modeling tissue-level transcript isoform patterns in cardiomyocytes and excitatory neurons

To establish controlled in vitro systems for interrogating lineage-specific gene deployment, human iPSCs were differentiated into cardiomyocytes and neurons using established protocols. Morphological progression confirmed successful lineage specification, with iPSCs transitioning through cardiac progenitor stages to beating cardiomyocytes by day 18, and through neural progenitor stages to mature neuronal networks by day 30 (Figure 7A). Immunofluorescence analysis further validated lineage identity, demonstrating expression of cardiomyocyte markers (*MYL2* and *MYL7*) and neuronal markers (*MAP2* and *NeuN*), confirming the acquisition of cardiac and neuronal phenotypes, respectively (Figure 7a).

**Figure 7a.**
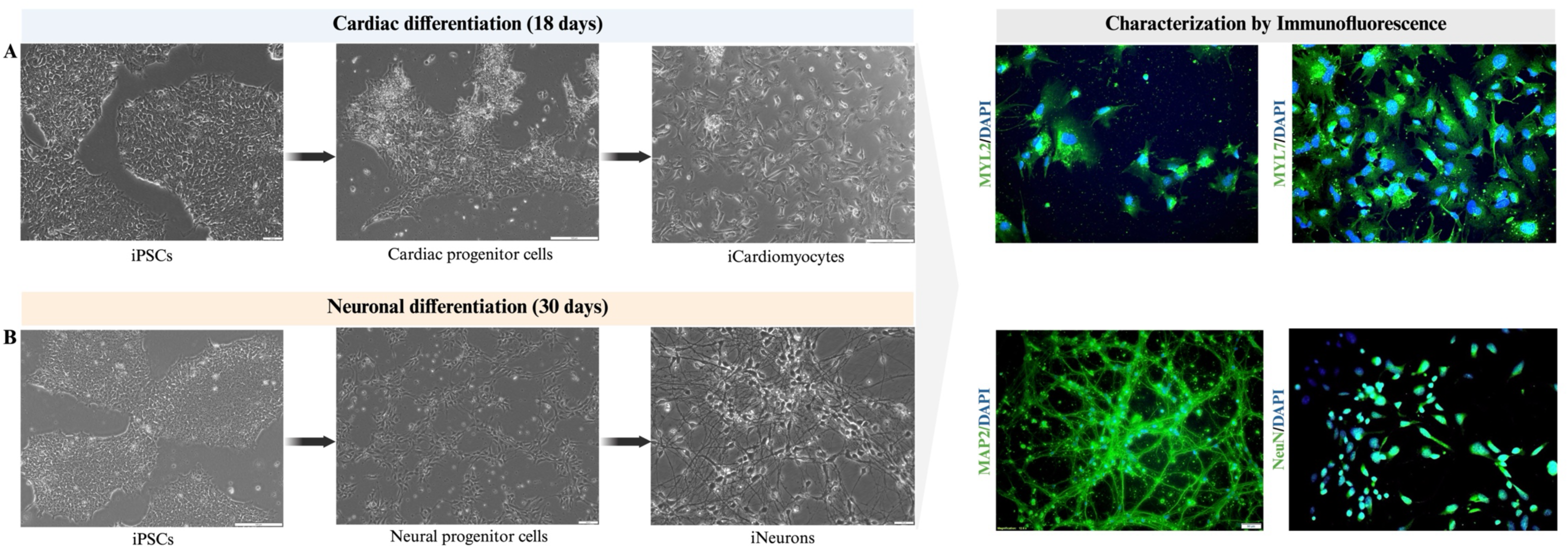
Differentiation and immunofluorescence characterization of iPSC-derived cardiomyocytes and neurons. (a) Schematic and representative phase-contrast images showing stepwise cardiac differentiation of human iPSCs into cardiac progenitor cells and mature iPSC-derived cardiomyocytes over 18 days. (b) Representative phase-contrast images illustrating neuronal differentiation of human iPSCs into neural progenitor cells and mature iPSC-derived neurons over 30 days. Immunofluorescence analysis confirms lineage identity of differentiated cells. iPSC-derived cardiomyocytes express the cardiac markers *MYL2* and *MYL7,* while iPSC-derived neurons show robust expression of neuronal markers *MAP2* and *NeuN*. Nuclei are counterstained with DAPI. Scale bars as indicated.

Single-cell transcriptomic profiling of these differentiated populations revealed well-defined cellular heterogeneity within each lineage (Figure 7b). In cardiomyocytes, distinct clusters corresponding to cardiac progenitor cells, proliferative cardiomyocytes, atrial cardiomyocytes (aCM), and ventricular cardiomyocytes (vCM) were identified. Similarly, neuronal datasets resolved major populations including neural progenitor cells, GABAergic neurons, layer 6 excitatory-like neurons, and early differentiating neurons. These results confirm that the in vitro systems capture key lineage-specific cell states relevant to cardiac and neuronal biology.

**Figure 7b.**
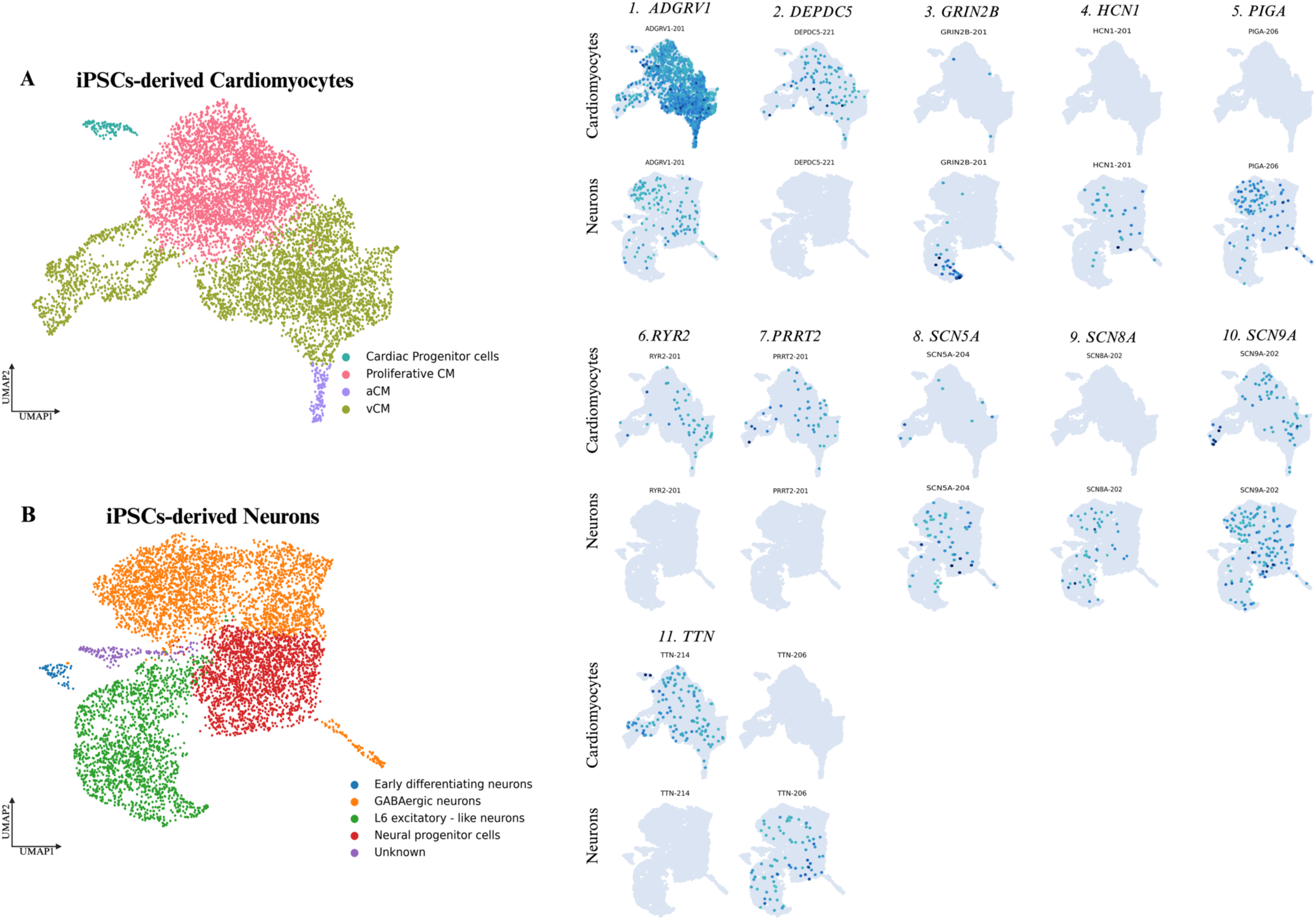
Cell-type–specific isoform expression patterns of shared epilepsy–SUDEP genes in iPSC-derived neurons and cardiomyocytes. (a) UMAP visualization of single-cell transcriptomes from human iPSC-derived neurons showing major neuronal subtypes, including neural progenitors, GABAergic neurons, and layer 6 excitatory-like neurons. (b) UMAP plot of iPSC-derived cardiomyocytes depicting cardiac progenitors, proliferative cardiomyocytes, atrial cardiomyocytes (aCM), and ventricular cardiomyocytes (vCM). (C-S) UMAP feature plots display isoform-resolved expression of selected genes across human iPSC-derived cardiomyocytes (top rows) and neurons (bottom rows). Each panel illustrates expression of individual transcript isoforms, highlighting cell-type–restricted transcript expression across neuronal and cardiac lineages. The genes shown include *ADGRV1*, *DEPDC5*, *GRIN2B*, *HCN1*, *PIGA*, *PRRT2*, *RYR2*, *SCN5A*, *SCN8A*, *SCN9A*, and *TTN*. Color intensity indicates normalized expression levels, with darker shading representing higher transcript abundance.

Consistent with the tissue-level analyses, examination of iPSC-derived cardiomyocytes and excitatory neurons revealed clear lineage-dependent expression patterns across epilepsy–SUDEP genes (Figure 7b). Several genes displayed preferential expression in one lineage while remaining detectable in the other, indicating partitioned rather than strictly tissue-exclusive expression.

Neuronal excitability and synaptic genes, including *GRIN2B, HCN1*, and *SCN8A*, were predominantly enriched in neuronal populations, with minimal or no detectable expression in cardiomyocytes. In contrast, a subset of genes showed preferential expression in cardiomyocytes, including *RYR2, PRRT2*, and *DEPDC5*, which were largely absent from neuronal populations. At the same time, some genes demonstrated detectable expression across both cell types, supporting the presence of shared genetic programs deployed in a lineage-dependent manner. For instance, *ADGRV1* and *SCN9A* were observed in both cardiomyocytes and neurons, with comparatively higher expression in cardiomyocytes and neurons, respectively. Isoform-level partitioning was also evident for genes such as *TTN*, where TTN-214 was detected predominantly in cardiomyocytes, whereas TTN-206 was observed in neuronal populations, further supporting lineage-specific deployment of transcript variants.

Although overall expression intensity and transcript diversity were reduced *in vitro* compared with native tissue datasets, likely reflecting differences in maturation state and microenvironmental complexity, the overall directionality of gene expression remained consistent with tissue-based observations. Notably, these patterns emerge in the absence of systemic physiological inputs, suggesting that lineage-dependent partitioning of gene expression is an intrinsic feature of cellular identity rather than a secondary consequence of tissue environment. These findings demonstrate that key aspects of the neuro–cardiac regulatory architecture identified in human tissues can be recapitulated in controlled in vitro systems, reinforcing the concept that shared genetic loci are differentially deployed across cardiac and neuronal lineages in a context-dependent manner.

## Discussion

This study was designed to connect population-level SUDEP genetics with a cell-type and isoform–resolved expression map across the two organs most plausibly involved in terminal collapse in SUDEP, brain and heart. Clinically, SUDEP events frequently follow generalized tonic–clonic seizures and progress through postictal dysfunction into central apnea, bradycardia, and asystole, supporting a convergent neuro–cardiorespiratory failure mechanism rather than a purely brain- or heart-restricted phenotype (Peng et al., 2017; Pysick et al., 2026; Ryvlin et al., 2013). Within this framework, the key observation from our study is not simply the shared genetic architecture between epilepsy and SUDEP, but that these shared genes are differentially deployed across tissues and lineages, including at the level of transcript isoforms. This suggests that risk variants may act through distinct molecular contexts in different organs, contributing to SUDEP vulnerability through multiple physiological substrates. This interpretation is consistent with prior works highlighting pleiotropy across neuronal and cardiac excitability pathways in SUDEP genetics (Friedman et al., 2018; Shlobin et al., 2024; Sun et al., 2023).

A central conceptual advance emerging from our results is a two-layer model of neuro–cardiac convergence in SUDEP. The first layer comprises a shared excitability substrate, where ion channels and transporters dominate the overlapping gene set between epilepsy and SUDEP. This aligns with extensive evidence implicating altered membrane excitability, synaptic transmission, and autonomic regulation as core mechanisms underlying seizure susceptibility and SUDEP risk. The second layer reflects a shift toward cardiac-associated biology within the SUDEP-enriched gene set, including genes involved in cardiac electrophysiology, calcium handling, and contractile function. These findings are concordant with clinical observations that fatal SUDEP events often involve postictal cardiorespiratory dysfunction, including arrhythmias and impaired cardiac output. Hence, this supports a model in which seizure-induced disruption of central autonomic regulation interacts with an underlying cardiac substrate, determining whether recovery occurs or progresses toward fatal collapse.

A major contribution of this study is the incorporation of isoform-resolved transcriptomics. Using long-read single-cell datasets, we directly assessed full-length transcript expression across cardiac and neuronal cell populations. This approach enabled classification of genes into three reproducible patterns: (i) heart-restricted isoforms, (ii) brain-restricted isoforms, and (iii) shared genes with comparative isoform expression across tissues. These patterns provide a mechanistic layer beyond gene-level overlap, demonstrating that tissue specificity is frequently encoded at the level of transcript isoforms rather than gene presence alone. This observation is consistent with the broader role of alternative splicing in defining tissue identity and functional specialization.

Within the comparative isoform category, several genes illustrate how shared loci can be deployed differently across tissues. For example, *SCN5A* showed multiple cardiac-associated isoforms together with a shared transcript detectable across both tissues, supporting its dominant role in cardiac electrophysiology while retaining broader expression potential. Similarly, *SCN9A* displayed both tissue-specific and shared transcript expression, consistent with its roles across excitable cell types. Structural genes further reinforce this concept: *TTN* exhibited extensive cardiac isoform diversity alongside shared and brain-detected transcripts, indicating that its functional contribution is context-dependent and not restricted to a single tissue. These findings suggest that interpretation of gene-level associations in SUDEP requires consideration of transcript-level context, particularly for genes with complex splicing architectures.

In contrast, a subset of genes demonstrated strong tissue-restricted isoform expression. Cardiac-restricted genes included *HCN4, KCNH2, KCNE1, MYH6, MYO18B*, and *ATP1A2*, all of which are closely linked to cardiac electrophysiology or contractile function. Brain-restricted genes included *SCN1A, SCN2A, SCN8A, GRIN2B, HCN1*, and *CACNA1A*, which are well-established regulators of neuronal excitability and synaptic transmission. These patterns reinforce the interpretation that epilepsy-associated risk is primarily mediated through neuronal pathways, whereas SUDEP-associated risk includes a stronger contribution from cardiac functional networks.

Importantly, several genes exhibited expression across both tissues but with distinct transcript expression, including *DEPDC5, ALDH5A1, SCARB2, TBC1D24*, and *RYR2*. These genes span pathways related to signaling, metabolism, membrane trafficking, and calcium handling, and their isoform partitioning suggests that shared molecular pathways may be differentially tuned across organs. Such genes cannot be categorized as exclusively neuronal or cardiac; rather, they represent points of convergence where perturbations may impact both systems simultaneously.

The *in vitro* iPSC-derived cardiomyocyte and neuronal models further support these findings. Although overall expression levels and transcript diversity were reduced relative to native tissues, consistent with differences in maturation and microenvironment, the directionality of lineage-dependent expression patterns was preserved. Genes such as *GRIN2B, HCN1*, and *SCN8A* remained neuron-enriched, whereas *RYR2* and *PRRT2* were primarily detected in cardiomyocytes. At the same time, genes including *ADGRV1, SCN5A*, and *SCN9A* were detectable across both lineages but showed clear differences in relative expression levels. Isoform-level partitioning was also retained, as illustrated by *TTN*, where distinct transcripts were preferentially associated with cardiomyocytes or neurons. Notably, these patterns emerge in controlled systems lacking systemic physiological inputs, indicating that lineage-dependent partitioning of gene and isoform expression is an intrinsic property of cellular identity rather than a secondary effect of tissue environment.

Overall, these findings argue against a model in which SUDEP represents merely a more severe form of epilepsy. Instead, the data support a framework of partial genetic overlap combined with divergence at the level of tissue deployment and transcript expression. Epilepsy risk maps predominantly to genes regulating neuronal excitability and synaptic signaling, whereas SUDEP risk shows additional convergence on cardiac electrophysiological and contractile pathways. Isoform-level regulation provides a plausible molecular mechanism underlying this divergence, enabling the same genetic loci to contribute to distinct physiological processes across organs. This multi-layered architecture supports a neuro–cardiac convergence model in which SUDEP arises from the interaction between seizure-driven central dysfunction and a susceptible cardiac substrate, rather than from a single-organ pathology.

## Limitations and Future Directions

While this study provides an integrative framework linking epilepsy and SUDEP genetics to tissue- and isoform-level expression patterns, several limitations should be considered. The genomic analysis is based on aggregation of published studies, introducing heterogeneity in sequencing platforms, variant-calling pipelines, and clinical annotation that may influence gene-level burden estimates. The absence of individual-level genotype data limited our ability to account for ancestry, comorbidities, and detailed clinical phenotypes, and prevented assessment of variant co-occurrence. In addition, SUDEP cases remain underrepresented in genomic datasets due to inconsistent recognition and classification, reducing statistical power and limiting genotype–phenotype resolution. Although long-read single-cell transcriptomic data enabled isoform-resolved analysis, these datasets may be influenced by differences in sample processing, sequencing depth, and annotation completeness; importantly, lack of detected expression does not imply biological absence but may reflect technical limitations such as dropout or limited detection sensitivity. Furthermore, iPSC-derived cardiomyocyte and neuronal models, while useful for assessing lineage-dependent gene deployment, do not fully recapitulate the maturation state or physiological complexity of native tissues. Future studies integrating population-matched cohorts with deep phenotyping, alongside long-read, spatial transcriptomic, and multi-omic profiling of well-characterized SUDEP cases, will be essential to validate candidate variants and define their functional impact, particularly through isoform-specific experimental approaches.

## Acknowledgments

We gratefully acknowledge the Al Jalila Foundation for providing foundational research infrastructure, institutional resources, and an enabling scientific environment, as well as for dedicated financial support through the Al Jalila Foundation Research Grant (AJF2023-103). Additional funding was provided by internal research grants from Mohammed Bin Rashid University of Medicine and Health Sciences (MBRU), College of Medicine (MBRU-CM-RG2024-12 and MBRU-CSRG-25-3). Dr. Binte Zehra was supported by the AJF funded MBRU Post-Doctoral Fellowship Award (MBRU-PD-2024-03) and the Saudi Heart Association Dr. Wael Al-Mahmeed Research Grant (EXT-RG2022-02, EXT-RG2025-01).

## Author Contributions

Conceptualization: B.Z., B.K.B., M.U. Methodology: B.Z., S.B.E., N.M., N.V., M.E., S.A-S. Formal Analysis: B.Z., S.B.E., M.F., R.T. Resources: M.U., B.K.B. Writing—Review & Editing: B.Z., S.B.E., M.F., N.V., I.A., M.A., S.A., S.D.P., B.K.B., M.U. Supervision: M.U., B.K.B. Visualization: B.Z., M.F., S.B.E., N.V. Funding Acquisition: M.U., B.K.B.

## Data Availability Statement

The raw and processed datasets generated from long-read single-cell sequencing have been deposited in the NCBI GEO database and can be accessed under the following accession numbers, iPSCs-derived Neurons: GSE274249 and iPSCs-derived Cardiomyocytes: GSE319652. Bulk tissue RNA-seq expression used in this study was obtained from the Genotype-Tissue Expression (GTEx) Project, V10. We used the open-access file: GTEx_Analysis_v10_RNASeQCv2.4.2_gene_tpm.gct.gz. For data processing and analysis, standard workflows were adopted, as described in the data Analysis section of the methodology.

## Legends

**Supplementary Figure 1.**
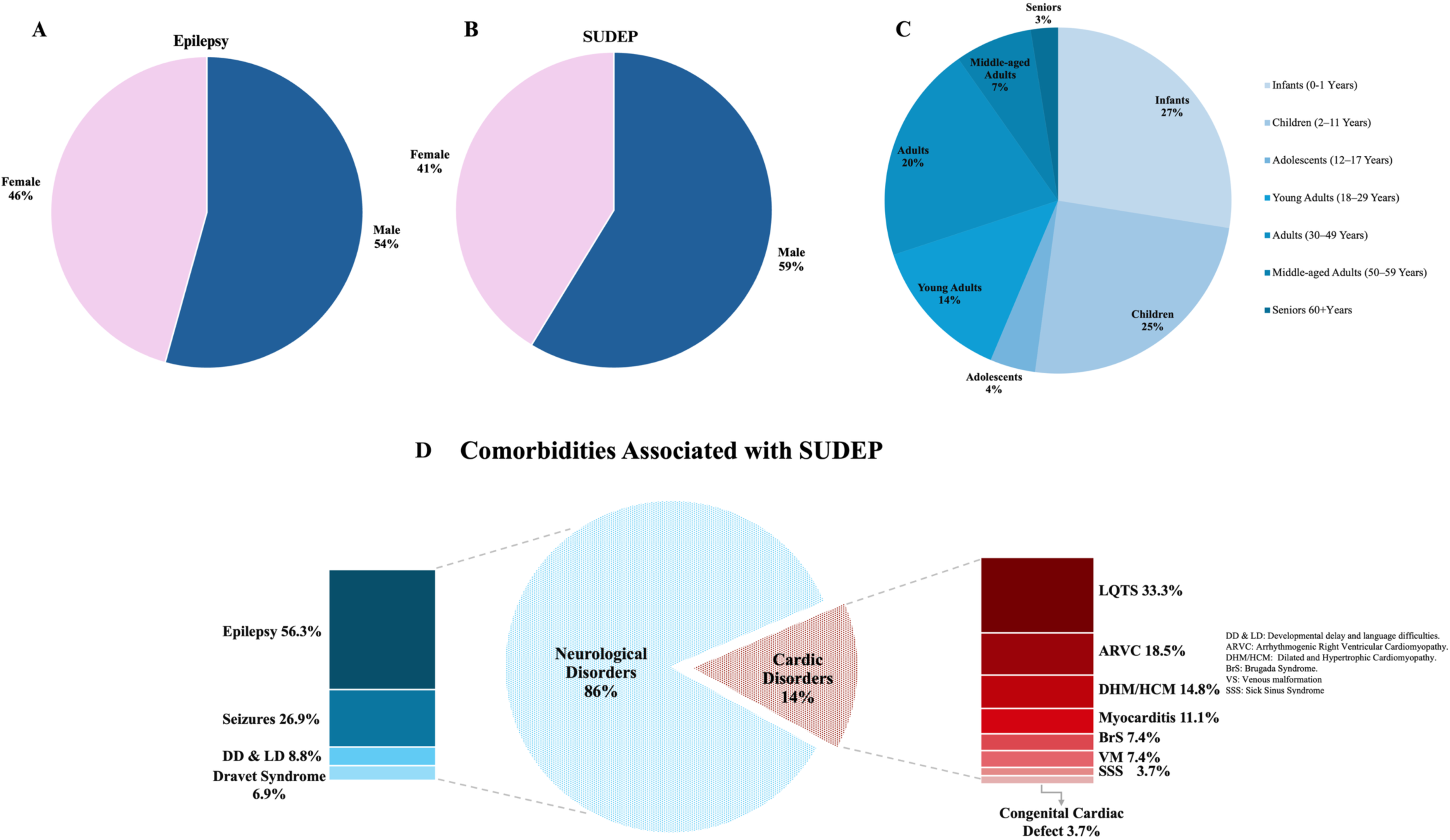
Demographic and clinical characteristics of SUDEP and epilepsy cohorts. (A–B) Gender distribution in epilepsy and SUDEP cohorts showing a male predominance in both groups (54% and 59%, respectively). (c) Age distribution of SUDEP cases (n=236) showing higher representation among infants (27%) and children (25%), followed by adults aged 30–49 years (20%) and young adults (14%). (d) Comorbidities associated with SUDEP (n=187) demonstrating that neurological disorders (86%) were most prevalent while cardiac disorders accounted for 14% of cases. Abbreviations: DD & LD, developmental delay and language difficulties; ARVC, arrhythmogenic right ventricular cardiomyopathy; DHM/HCM, dilated/hypertrophic cardiomyopathy; BrS, Brugada syndrome; VM, venous malformations; SSS, sick sinus syndrome.

**Supplementary Figure 2.**
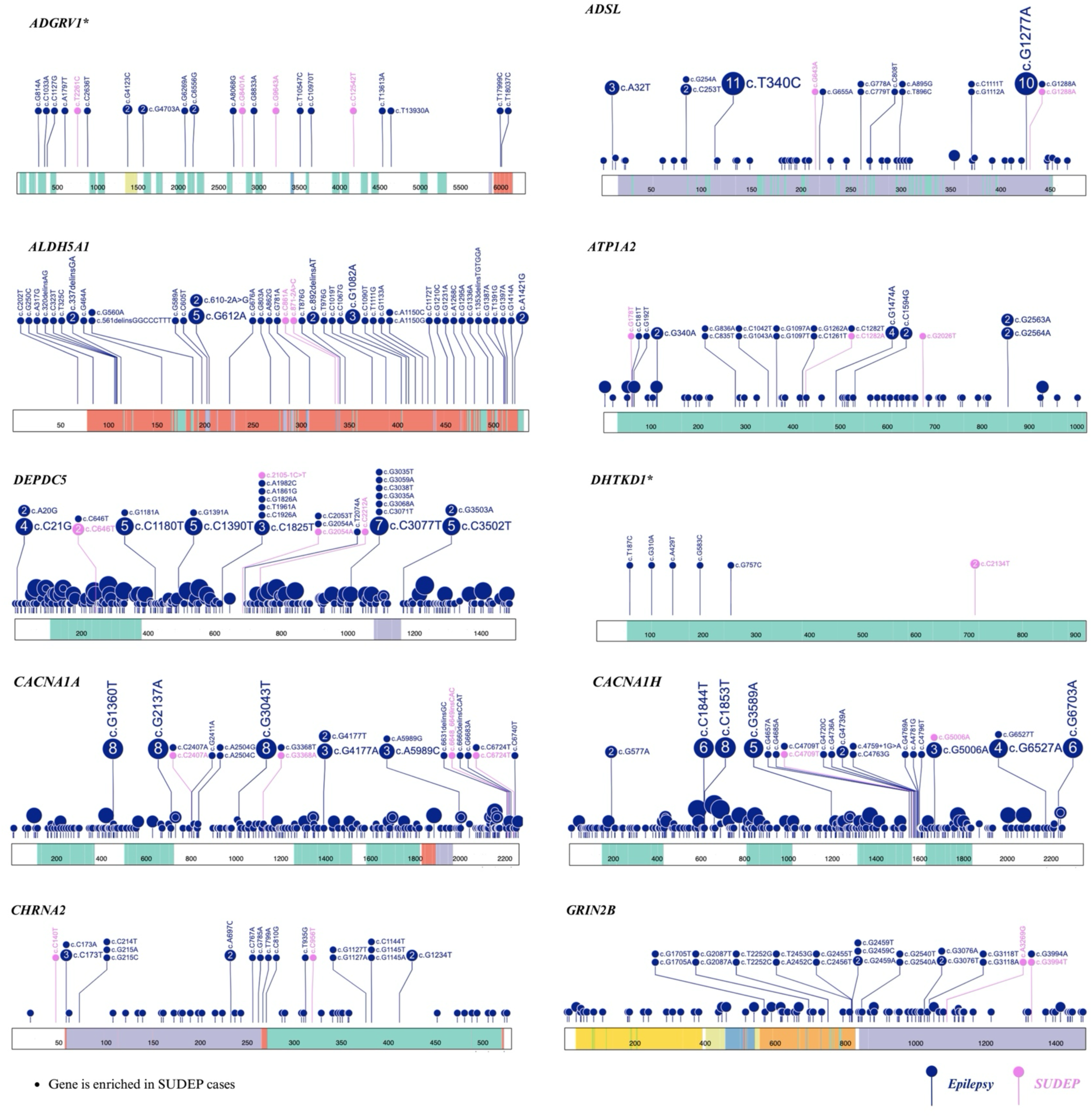
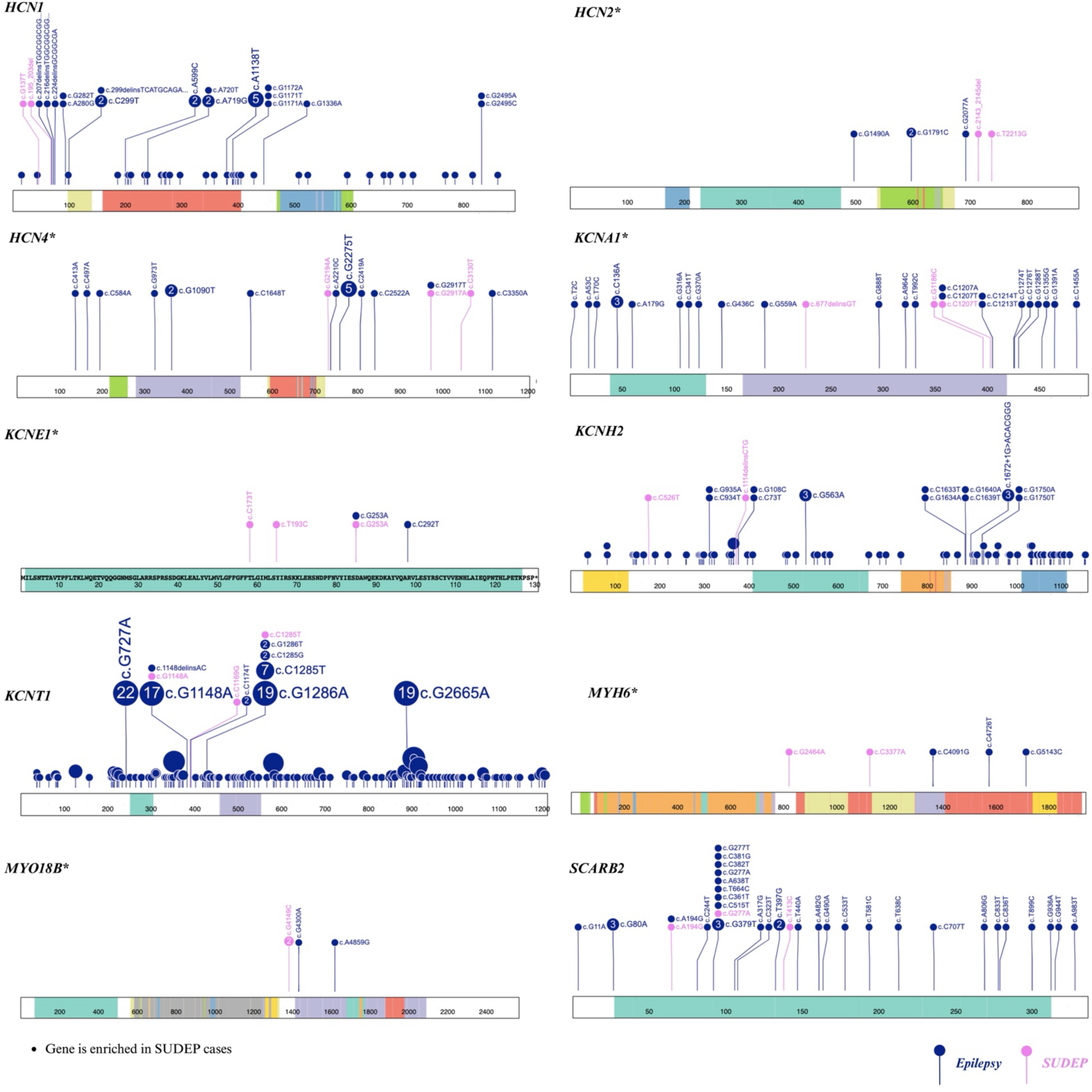
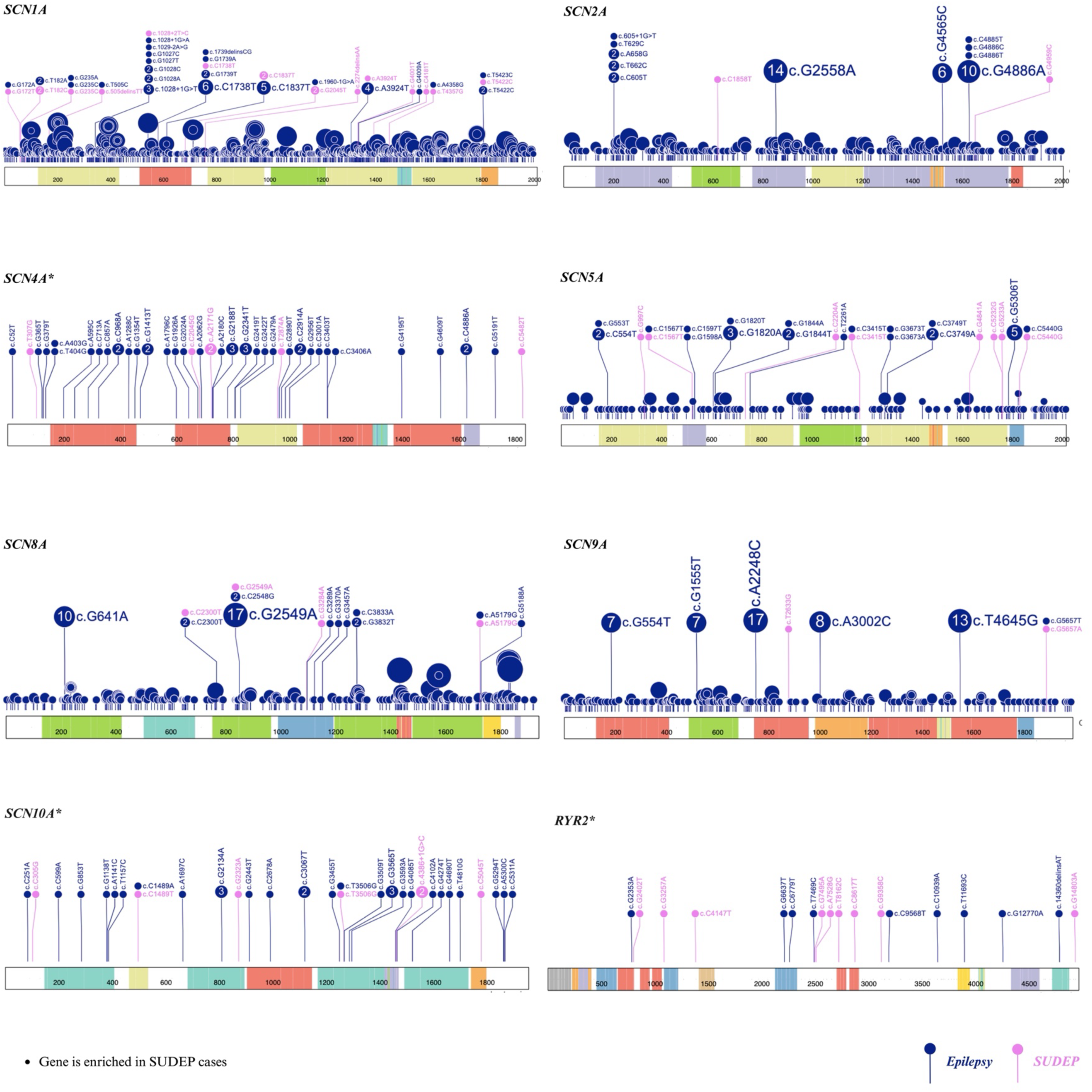
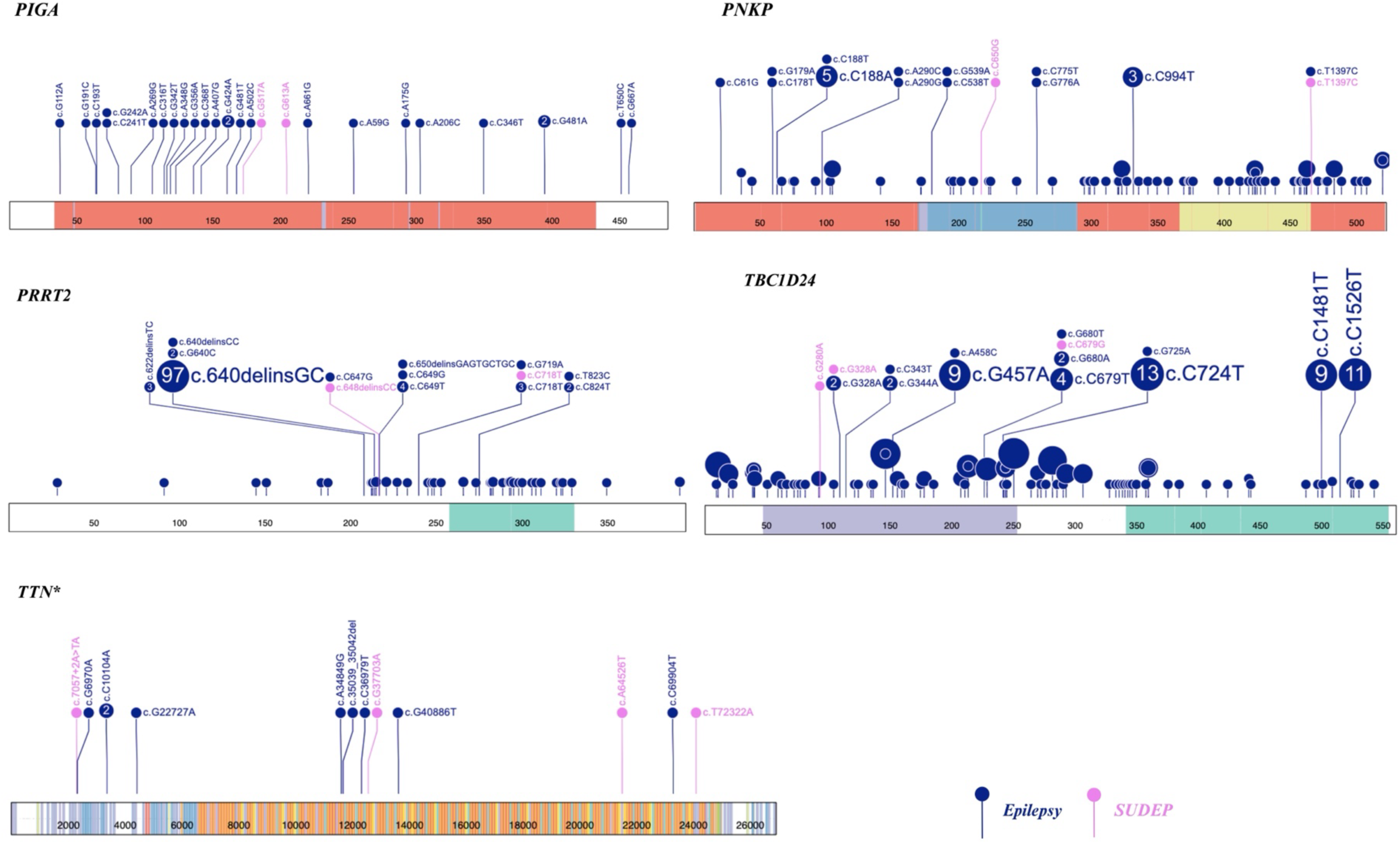
Domain-level distribution of rare variants across 33 overlapping genes in epilepsy and SUDEP cohorts. Lollipop plots illustrate mutational hotspots mapped onto annotated protein domains for each gene, distinguishing SUDEP-associated variants (pink) and epilepsy-associated variants (blue). Shared variants present in both phenotypes are represented by dual-colored markers.

**Supplementary Figure 3.**
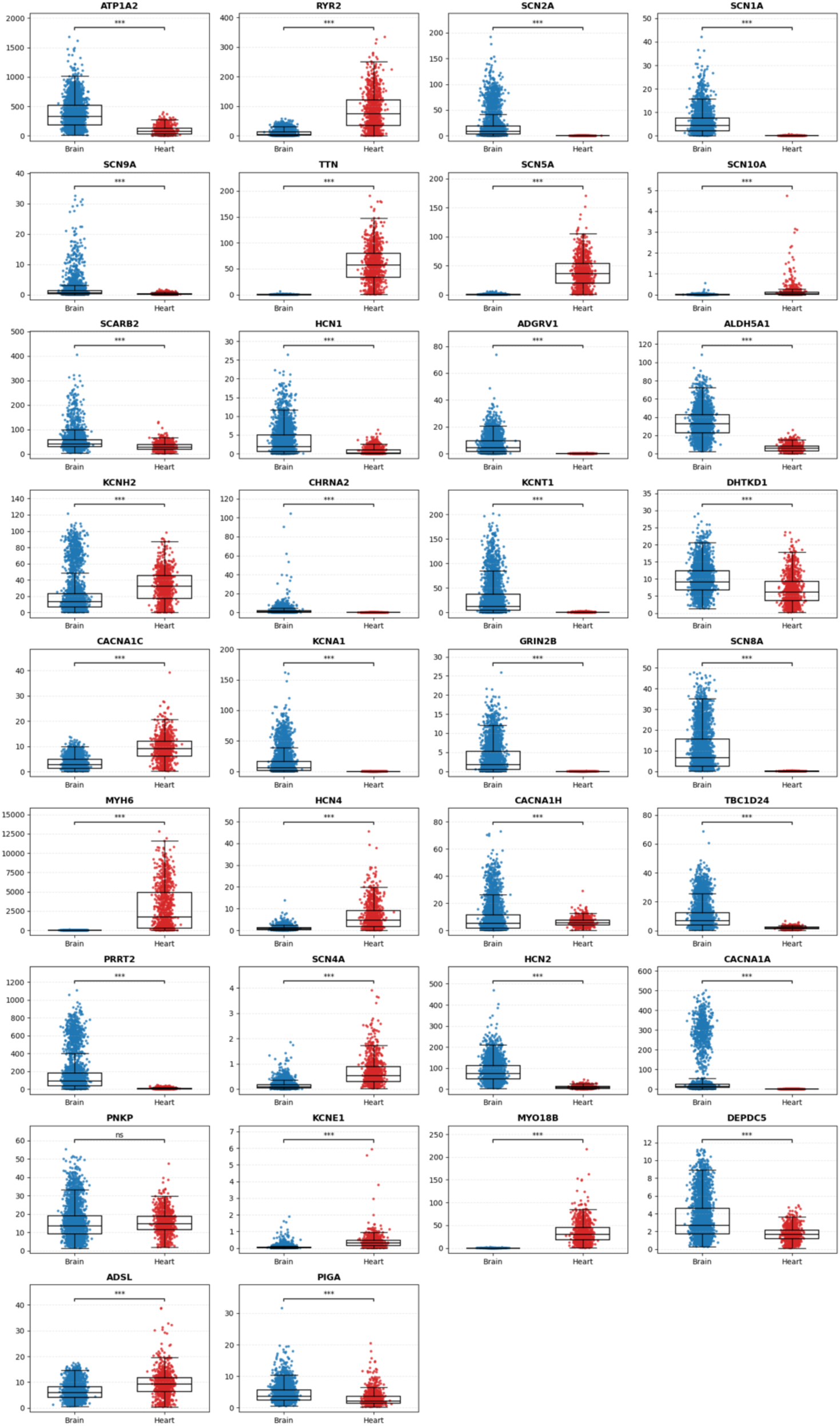
Comparative expression analysis of 33 overlapping epilepsy–SUDEP genes across human brain and heart tissues using GTEx RNA-seq TPM values. Boxplots represent transcript abundance (TPM, log-scaled) in brain (blue) and heart (red) tissues for each gene. Statistical significance is indicated by asterisks (\**p* < 0.05, \*\**p* < 0.01, \*\*\**p* < 0.001), highlighting tissue-specific expression divergence among shared neuro-cardiac genes.

**Supplementary_metadata.** List of papers included and excluded from the meta-analysis for epilepsy and SUDEP cohorts.

**Supplementary S1.** Variant annotated and hg38 harmonized SUDEP variant dataset.

**Supplementary S2.** Functional categorization and variant distribution of genes shared between epilepsy and SUDEP. The table reports gene function, chromosomal location, variant counts in SUDEP and epilepsy, variant types and regions, and gene-level enrichment estimated using posterior rate ratios (RR) with credible intervals (CrI) and FDR correction.

**Supplementary S3.** Isoform-level expression of genes of interest across publically available heart and brain isoform atlases.

**Supplementary S4.** Isoform-level expression of genes of interest across iPSC-derived cardiomyocytes (iCMCs) and iPSC-derived neurons (iNeurons) in the long-read single-cell transcriptomic dataset.

